# Plant biomechanics and resilience to environmental changes are controlled by specific lignin chemistries in each vascular cell type and morphotype

**DOI:** 10.1101/2021.06.12.447240

**Authors:** Delphine Ménard, Leonard Blaschek, Konstantin Kriechbaum, Cheng Choo Lee, Henrik Serk, Chuantao Zhu, Alexander Lyubartsev, Nuoendagula, Zoltán Bacsik, Lennart Bergström, Aji Mathew, Shinya Kajita, Edouard Pesquet

**Author notes:** These authors contributed equally (Delphine Ménard, Leonard Blaschek).

## Abstract

The biopolymer lignin, deposited in the cell walls of vascular cells, is essential for long-distance water conduction and structural support of plants. Independently of the species, each different vascular cell type contains a conserved lignin chemistry with specific aromatic and aliphatic substitutions. Yet, the biological role of this conserved and specific lignin chemistry for each cell type remained unclear. Herein, we investigate the role of specific lignin chemistries for cellular function by producing single cell analyses on vascular cell morphotypes, all enabling sap conduction but differing in morphology. We found that specific lignin chemistries accumulate in each morphotype. Moreover, lignin accumulates dynamically, increasing in quantity and changing composition, to alter the cell wall biomechanics of each morphotype during their maturation. For similar aromatic substitution, residues with alcohol aliphatic functions increased stiffness whereas aldehydes increased flexibility. Modifying this specific lignin chemistry impairs the cell wall biomechanics of each morphotype and consequently reduces their capacity to optimally conduct water in normal conditions, and to recover from drought. Altogether, lignin chemistry is differently controlled for each sap conducting cell types during their maturation to dynamically adjust their biomechanics and hydraulic properties to adapt to developmental and environmental constraints.

## Introduction

Vascular plants have a unique tissue called xylem that functions as both a load-bearing skeleton and a conductive system for long-distance water transport. This dual function depends on the xylem conduit cells called tracheary elements (TEs). During their differentiation, TEs reinforce their primary cell walls (PCWs) with patterned secondary cell walls (SCWs), and subsequently remove their intracellular content by programmed cell death. This hollowing out of TEs starts their water conducting function, forming unobstructed tubes with thickened and regularly patterned sides (Ménard et al. 2015; Derbyshire et al. 2015). As the plant grows, new TEs form, die and connect both longitudinally and laterally to other TEs to conduct water throughout the plant (Ménard and Pesquet 2015). Because TEs require cell death to function, they have been considered inert and unable to adjust to changing developmental and environmental constraints. However, TE cell walls undergo modifications after their death, such as the continuous accumulation of lignin (Pesquet et al. 2010; Pesquet et al. 2013; Smith et al. 2013; Derbyshire et al. 2015; Blaschek, Champagne et al. 2020). This *post-mortem* lignification is catalysed by cell wall embedded oxidative enzymes in TE cell walls and a cooperative supply of monomers by surrounding cells (Barros et al. 2015). The oxidative polymerisation of lignin fills the gaps between the cellulose and hemicellulose polymers in the cell walls formed before TE cell death (Blaschek and Pesquet 2021). To respond to developmental and environmental changes, the xylem forms different TE morphotypes with specific dimensions and SCW patterns such as narrow protoxylem (PX) with annular or spiral patterns, wide metaxylem (MX) with reticulate or pitted patterns, or later forming secondary xylem (SX) with SCW patterning similar to the MX but with narrower diameters. Additionally, the xylem also changes the cell types surrounding each TE morphotype, varying the proportion of neighbouring TEs, unlignified xylem parenchyma (XP) and lignified xylary fibres (XF) (Fig. S1; Chaffey et al. 2002; Derbyshire et al. 2015).The xylem sap, mainly water, ascends in the lumens of the dead interconnected TEs due to a negative pressure pull caused by gradients of water potential (Ψ) along the soil-plant-atmosphere continuum, and is finally released by evapotranspiration through the leaves. Unlike the Ψ of soil and air that vary with environmental conditions (Fig. S1), plants actively control their Ψ using both stomatal movements to regulate leaf transpiration rates, and the intra-cellular osmolarity of XPs and XFs to alter osmotic pressure (Bentrup 2017; Holbrook et al. 1995; Pockman et al. 1995). TE optimal morphology for laminar sap flow is a cylindrical pipe according to the Hagen-Poiseuille law (Calkin et al. 1986; Tyree et al. 1994; Venturas et al. 2017). TEs cannot however withstand very large Ψ differences as encountered during drought, which cause TEs to collapse inwardly, altering their circularity and consequently disrupting plant hydraulic conductivity (Fig. S1; Cochard et al. 2004; Brodribb and Holbrook 2005; Zhang et al. 2016; Coleman et al. 2008; Kitin et al. 2010; Voelker et al. 2011). TE inward collapse is sometimes reversible, acting as a circuit breaker to resist extreme drought, which is restored once water availability improves (Zhang et al. 2016). TE inward collapse is also observed when modifying the biosynthesis of TE SCWs using genetic mutations (Turner and Sommerville 1997; Brown et al. 2005) or drug treatment (Amrhein et al. 1983; Smart and Amrhein 1985) where it is called *irregular xylem* (*irx*). This *irx* is only observed in dead sap conducting TEs but never in non-sap conducting TEs produced either in isolated suspension (Endo et al. 2008) or ectopically in non-xylem tissues (Takenaka et al. 2018). The *irx* reveals the importance of TE cell wall composition, concentration and/or structure to set a mechanical resistance sufficient to cope with Ψ variations of soil and atmosphere. Lignin formation is a genetically controlled process regulating concentration, composition and structure of lignin polymers during the development and stress response for each cell type in their different cell wall layers (Blaschek, Nuoendagula et al. 2020; Blaschek, Champagne et al. 2020; Yamamoto et al. 2020; Hiraide et al. 2021). Lignin main residues are C_6_C_3_ phenylpropanoids (Tab. S1; Moss 2000) that differ by their C_6_ aromatic *meta* groups, such as monomethoxylated guaiacyl (**G**) and dimethoxylated syringyl (**S**) rings, and in their C_3_ aliphatic terminal functions, such as alcohol (X_CHOH_) or aldehyde (X_CHO_) (Dixon and Barros 2019). TEs accumulate mostly **G** residues in their cell walls independently of the plant species (Pesquet et al. 2019). Distinct TE morphotypes also differently accumulate X_CHO_ residues (Yamamoto et al. 2020). Additionally, non-canonical residues are also incorporated in lignin such as benzaldehydes (H. Kim et al. 2003; Ralph et al. 2001), coumaroyl esters (Lapierre et al. 2021), stilbenoids (del Río et al. 2017; Rencoret et al. 2019), flavonoids (Lan et al. 2015; Rencoret et al. 2022) and other residues presenting phenyl (P) rings (Faix and Meier 1989; Kawamoto 2017). Yet, the role of this cell and morphotype specific lignin chemistry for TEs is still not understood.

Herein, we have investigated the biological roles of the specific lignin chemistry of the different TE morphotypes on the load-bearing and vascular properties of plants. We used three complementary biological systems to fully investigate lignin in TEs: (i) inducible plant pluripotent cell suspension cultures (iPSCs) from *Arabidopsis*, (ii) annual herbaceous *Arabidopsis* plants with genetically altered lignins, and (iii) perennial woody poplar plants with genetically altered lignins. IPSCs enable modifying lignin in isolated TE morphotypes at distinct *post-mortem* maturation stages and investigate the role of lignin in isolated TEs independently of the physiological constraints of sap conduction or tissue pressure. Genetically modified plants with altered lignin amounts and compositions enable assessing the role of specific lignin chemistry on the different sap-conducting TE morphotypes embedded in functional vascular tissues. The perennial poplar plants additionally enable to monitor the dendrochronological changes of lignin in TEs within tissues during secondary growth. Using these complementary systems, we could show that different TE morphotypes accumulate specific lignins during *post-mortem* lignification for optimal hydraulic properties. More specifically, the proportions of different C_3_ terminal functions for the same C_6_ substitution balanced cell wall stiffness with flexibility. Altogether, our study suggests that lignin structure is specificallyfine-tuned during the *post-mortem* maturation of each functioning TE morphotype to dynamically adjust their conductive and load-bearing properties to changing developmental and environmental conditions.

## Results

### *Post-mortem* accumulation of lignin increases the cell wall stiffness of isolated TEs morphotypes

To define the impact of lignin accumulation on the cell walls of isolated TEs, we used iPSCs as they enable to follow the maturation of intact single cell TEs that cannot easily be isolated from whole plants using maceration or dissection (Ménard et al. 2017). IPSCs from *Arabidopsis thaliana* synchronously produced all TE morpho-types, which underwent cell death 5–7 days after induction, followed by *post-mortem* lignification (Pesquet et al. 2010; Pesquet et al. 2013; Derbyshire et al. 2015). We monitored these *postmortem* changes in TEs at the nanoscale level using scanning electron microscopy (SEM) coupled with energy-dispersive X-ray spectroscopy (EDS) by measuring elemental changes in cell wall composition (Fig. 1A,B). Ratios of carbon (C) to coating chromium (Cr) content showed gradual *post-mortem* increases of C-rich compounds only in SCWs compared to PCWs, which plateaued by day 40-50 (Fig. 1C). Ratios of carbon to oxygen (O) content revealed that the compounds accumulating *post-mortem* in TE SCWs were lowly oxygenated as expected from lignin (9 C:3–5 O for lignin-units compared to 6 C:6 O for cellulose units; Fig. 1C). Once plateau was reached, PX TEs presented SCWs with significantly less C/Cr and C/O content than MX TEs, indicating differences in cell wall polymer amounts and composition between TE morphotypes (Fig. S2). Complementary biochemical analyses confirmed that the *postmortem* changes in cell wall corresponded to increases in lignin concentrations rising gradually only in TE cells to ∼25% of total dry cell wall weight by day 30 (Fig. S2). The impact of *postmortem* lignification on TE biomechanics was then evaluated using atomic force microscopy (AFM) on 5–20 μm2 areas of isolated 10and 50-day-old PX and MX TEs (Fig. 1D). AFM analysis showed that *post-mortem* lignification had almost no impact on PCWs but altered the biomechanics of SCWs for each TE morphotype by significantly increasing stiffness and decreasing deformability (Fig. 1E). Altogether we showed that SCWs of each TE morphotypes continue lignifying for more than 40 days after cell death to accumulate specific amount of lignin to increase the stiffness of TE SCWs.

**Figure 1.**
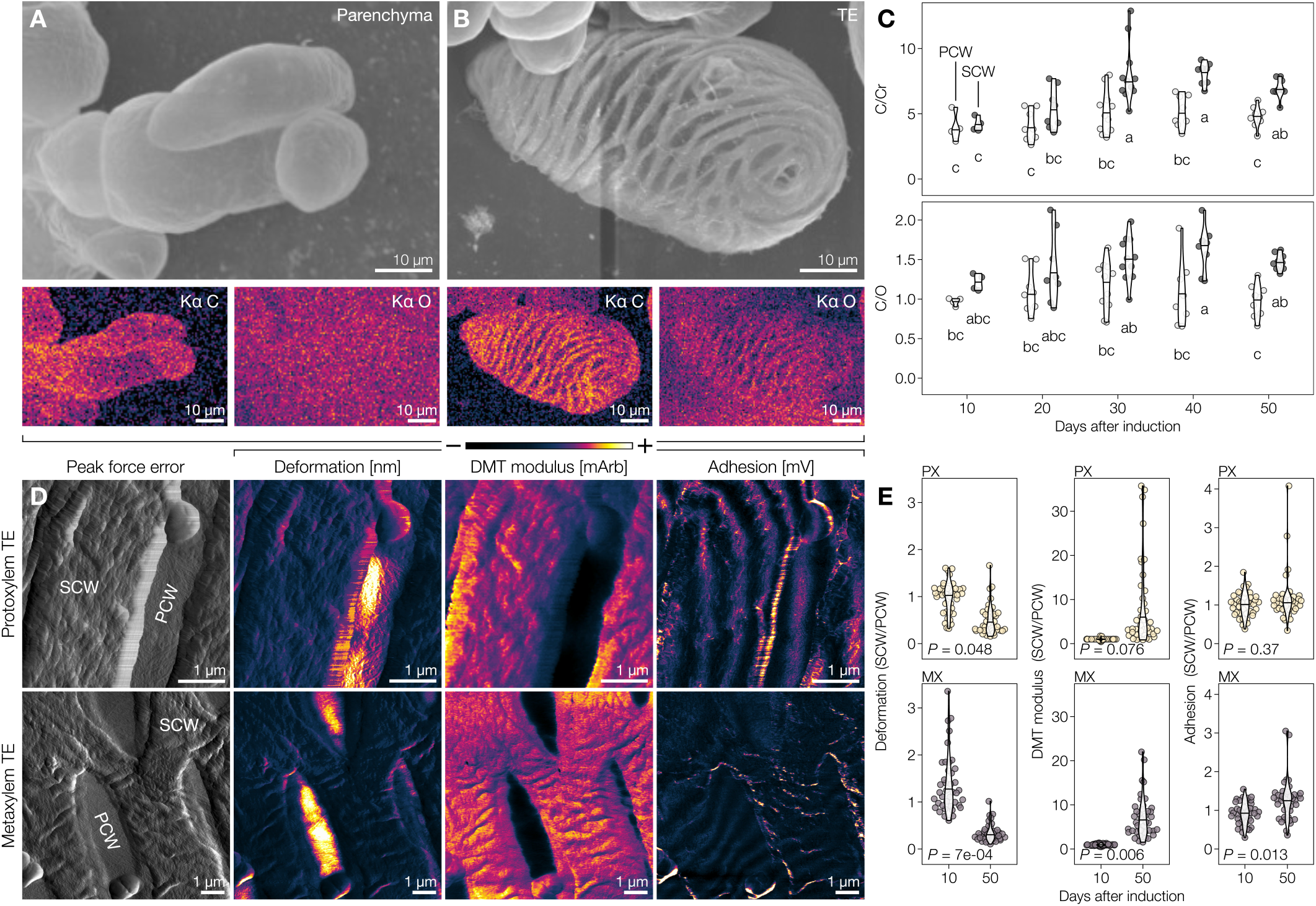
Post-mortem lignification actively alters biomechanics of TE secondary cell walls (SCWs). **A** Scanning electron micrograph of isolated parenchyma cells, prepared using critical point drying (CPD), as well as its energy-dispersive X-ray spectroscopy (EDS) carbon (C) and oxygen (O) signals in colour-coded intensity. **B** Scanning electron micrograph of an isolated tracheary element (TE) 30 days after induction, prepared by CPD, as well as colour-coded intensity EDS C and O signals. **C** EDS ratios of C to coating chromium (Cr) and C to O ratios of 9 μm^2^ (300 nm sided square) of the primary (PCW) and secondary cell wall (SCW) of isolated 10-to 50-day-old TEs. Note that SCWs gradually increase their C/O ratio during *post-mortem* maturation; *n* = 4–11 individual cells per time-point. Letters indicate significant differences according to a Tukey-HSD test (per panel; α = 0.05). **D** AFM peak force error as well as intensity color-coding of deformation, DMT modulus and adhesion of 50-day-old isolated protoxylem (PX) and metaxylem (MX) TEs. **E** SCW to PCW ratios of deformation, DMT modulus and adhesion of 10- and 50-day-old PX and MX TEs. P-value of a two-tailed Welch’s t-test is indicated; *n* = 4 individual cells per time-point and morphotype, 10 measurements per cell.

### *Post-mortem* lignin accumulation controls the resistance of isolated TE morphotypes to negative pressure

Next, we evaluated the role of stiffness increases due to *post-mortem* lignification in TE resistance to negative pressures such as those faced during water conduction. Isolated 10- and 50-day-old TEs were exposed to two different drying methods and observed using SEM. Critical-point drying (CPD), which minimizes negative pressure differences during the drying process, was compared to air drying, which mimics drought by exposing the cells to large Ψ differences. Parenchymatic cells showed no inward collapse after CPD but were completely flattened by air drying (Fig. 2A,B). TEs were similarly unaffected by CPD but partly withstood air drying (Fig 2A,B). These results showed that TE collapse occurs in response to the negative pressure exerted on single TE itself and is not due to surrounding tissue pressure. Analysis of the proportion of collapsed TEs after air drying during *post-mortem* lignification revealed a gradual increase of resistance to collapse, with the majority of 50-day-old TEs remaining completely intact (Fig. 2C,D). As the susceptibility to collapse lowered as *post-mortem* lignification increased, our results moreover showed that the susceptibility to collapse did not depend on the *pre-mortem* deposited SCW polysaccharidic polymers but rather on the free spaces in cell wall that these defined for lignification. Both PX and MX TEs (Fig. 2E,F) withstood collapse in air drying better as *post-mortem* lignification progressed, although PX were consistently more sensitive than MX TEs (Fig. 2C,D). To demonstrate that lignins caused the observed increased resistance, TE SCW lignification was completely prevented with piperonylic acid (PA) treatment (Fig. S2; Van de Wouwer et al. 2016; Decou et al. 2017). The resulting unlignified TEs completely collapsed with air drying (Fig. 2G). These results set that lignin amounts in SCWs control TE biomechanical properties to resist collapse. We thus show that the role of *post-mortem* lignification in TEs is to dynamically reinforce the cell walls of TEs to sustain the changes in negative pressure gradients associated to sap conduction during developmental and environmental constraints.

**Figure 2.**
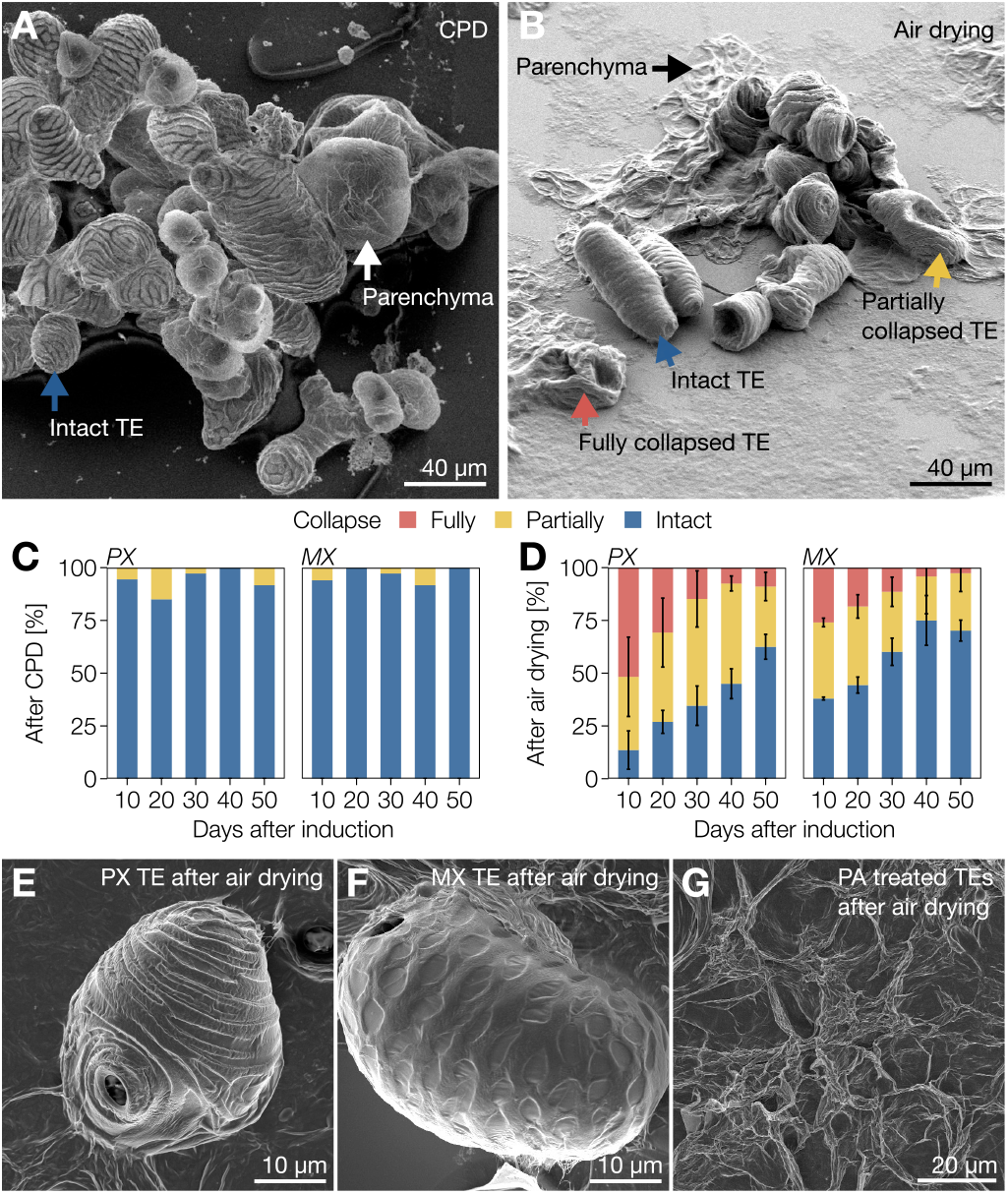
Gradual *post-mortem* lignification enables all TE morpho-types to resist extreme Ψ differentials. **A** Scanning electron micro-graph of 30-day-old isolated TEs and parenchyma cells produced from iPSCs and prepared using critical point drying (CPD). Note that both TEs and parenchyma are intact as indicated by blue and white arrows respectively. **B** Scanning electron micrograph of 30-day-old isolated TEs and parenchyma cells produced from iPSCs and prepared using air drying. Note that parenchyma cells (black arrow) are completely flattened whereas TEs were either fully collapsed (red arrow), partially collapsed (yellow arrow) or intact (blue arrow). **C** Relative proportion of 10-to 50-day-old TEs from iPSCs that were fully collapsed, partially collapsed, or intact after CPD; *n* = 7–50 individual cells per cell type and time-point. **D** Relative proportion of 10-to 50-day-old TEs from iPSCs that were fully collapsed, partially collapsed, or intact after air drying. Error bars represent ± SD of 2 independent experiments; *n* = 18–50 individual cells per cell type and time-point. **E** Scanning electron micrograph of a 30-day-old PX TE after air drying. **F** Scanning electron micrograph of a 30-day-old MX TE after air drying. **G** Scanning electron micrograph of 30-day-old unlignified TEs treated with PA after air drying.

### Each TE morphotype in annual plants differs in morphology, lignin concentration and composition

We then investigated the role of lignification on the collapse of TEs embedded in tissues using the *Arabidopsis thaliana* herbaceous plants. Three TE morphotypes are present in 8-week-old stems of wild-type (WT) plants, identifiable by their distance to the cambium: protoxylem TEs (PX), metaxylem TEs (MX), and secondary TEs (SX) (Fig. 3A). For all morphotypes, cell types directly adjacent to each TE included always 35% of other TEs but various proportions of XP and XF (Fig 3B). PX and SX had smaller lumen diameters compared to MX (Fig. 3C). Semi-quantitative lignin analysis *in situ* was performed using Raman spectroscopy and calibrated respectively to key lignin mutants (Fig. S3; Blaschek, Nuoendagula et al. 2020). MXs presented higher lignin levels than PX (Fig. 3D). All TEs exhibited similar **S**/**G** residue proportions (Fig. 3E). The ratio of terminal coniferaldehyde (**G**_CHO_) to total coniferyl alcohol (**G**_CHOH_; including terminal coniferyl alcohol as well as internal guaiacylglycerol, pinoresinol and other incorporated **G**_CHOH_ structures) was higher in the SX than in PX or MX (Fig. 3F).The proportion of non-canonical benzaldehyde and **P** residues also varied between morphotypes, increased in PX compared to MX and SX (Fig. S4). Total **G**_CHO_, measured using the Wiesner test (Blaschek, Champagne et al. 2020), showed that PX accumulated less **G**_CHO_ than MX and SX (Fig. S4). Overall, each TE morphotype had specific dimensions, SCW organisation and neighbouring cells as well as distinct lignin composition, amounts and structure.

**Figure 3.**
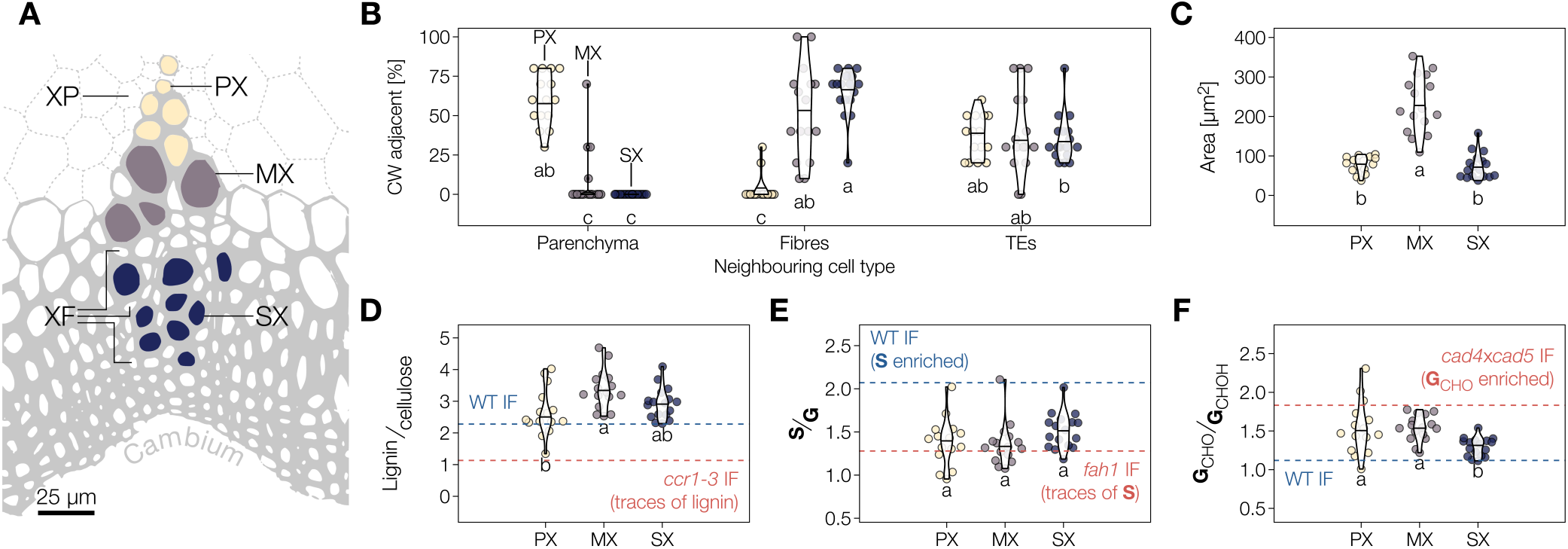
Different TE morphotypes in annual plants have specific morphological features and lignin chemistry. **A** Scheme of the localisation of the three TE types in vascular bundles of *Arabidopsis* stems according to their distance to the cambium: protoxylem (PX) in yellow, metaxylem (MX) in purple and secondary xylem (SX) in blue. Each TE morphotype shared different proportions of xylary fibres (XF) and xylem parenchyma (XP) surround each TE type. **B** Relative proportion of adjacent cell type for each TE morphotype. Note that the proportion of neighbouring TEs remains constant independently of the TE morphotype. Letters indicate significant differences according to a Kruskal-Wallis test (Holm adjusted for multiple comparisons; α = 0.05). **C** Lumen area of each TE morphotype determined from cross sections. **D** Relative lignin to cellulose ratio measured by Raman microspectroscopy. **E** Relative **S** to **G** ratio measured by Raman microspectroscopy. **F** Relative **G**_CHO_ to **G**_CHOH_ ratio measured by Raman microspectroscopy. Because range and intercept of Raman band ratios differ from other biochemical analyses but still maintain a linear relationship (Agarwal 2019; Blaschek, Nuoendagula et al. 2020), references are presented for each lignin parameter using interfascicular fibers in WT and relevant mutants. Different letters in panels C–H indicate significant differences according to a Tukey-HSD test (per panel; α = 0.05); *n* = 15–17 individual cells per TE type in 3 plants.

#### Lignin concentration and composition differently affects the resistance to negative pressure of each TE morpho-type in annual plants

To determine the importance of accumulating specific lignins for the mechanical resistance of sap transporting TEs, we evaluated TE lignification and collapse in nine loss-of-function mutants altered in lignin concentration and/or composition (list of genes mutated is provided in Tab. S2). We measured TE morphology as well as lignin structure using *in situ* quantitative imaging (Blaschek, Nuoendagula et al. 2020; Blaschek, Champagne et al. 2020; Yamamoto et al. 2020). For TE morphology, we measured inward collapse, estimated by a decreased convexity, and deformation, shown by a reduction of circularity (general deviation from a perfect circular shape) compared to TEs in WT plants (Fig. 4A,B).The used mutant array provided in a dataset with around 100 TEs per morphotype with a wide and continuous variation in convexity and lignin structure (Fig. S5), suitable to detect specific associations between lignin amount and composition and TE resistance to collapse for each morphotype. TE perimeter and neighbouring cell types remained unaltered in all TE morphotypes between mutants, thereby confirming that neither TE *pre-mortem* formation nor surrounding cells were altered by mutations (Fig. S4). In contrast, convexity and circularity of specific TE morphotypes were reduced by genetic changes leading to various degrees of deformation (Fig. 4C), such as *ccr1* affecting all TE morphotypes whereas *ccoaomt1* altered only MXs. To define which lignin chemical parameters affected the inward collapse of each TE morphotype, we computed structural equation models to identify significant associations. In addition to the positive effect of higher lignin amounts, increases in terminal **G**_CHO_, total **G**_CHOH_ contents and TE perimeter were associated with higher resistance to collapse in PXs (Fig. 4D). MX resistance to collapse was similarly associated with higher levels in total lignin and **G**_CHO_ (both total and terminal), whereas increases in **S** residues and in the proportion of neighbouring TEs decreased their resistance to collapse (Fig. 4E). The association between increases in lignin and lower susceptibility to TE collapse for both PX and MX directly confirmed our observations made of isolated TEs from iPSCs (Fig. 2). TE collapse was rare in SXs (Fig. 4A) and only associated with lower levels of **G**_CHOH_ and high levels of benzaldehydes (Fig. 4F). We also performed interaction analyses to determine the interdependence between morphological and biochemical features for TE susceptibility to collapse (Fig. S5). In PXs, the synergistic strengthening effects of increased levels of **G**_CHOH_ and **G**_CHO_ that promoted TE resistance depended on TE perimeter (Fig. S5). In MXs, the positive effect of higher **G**_CHO_ was synergistic with total lignin amounts to promote TE resistance but depended on neighbouring TE proportions (Fig. S5). Overall, TE susceptibility to collapse in each TE morphotype was associated with specific changes and interactions between cell/tissue morphology and lignin chemistry (**S**/**G** and **G**_CHO_/**G**_CHOH_ compositional ratios, (**G**_CHO_ terminal to total). Our results set that the different lignin residues have non-redundant roles and need to be specifically controlled for each TE morphotype to sustain negative pressure for optimal water conduction.

**Figure 4.**
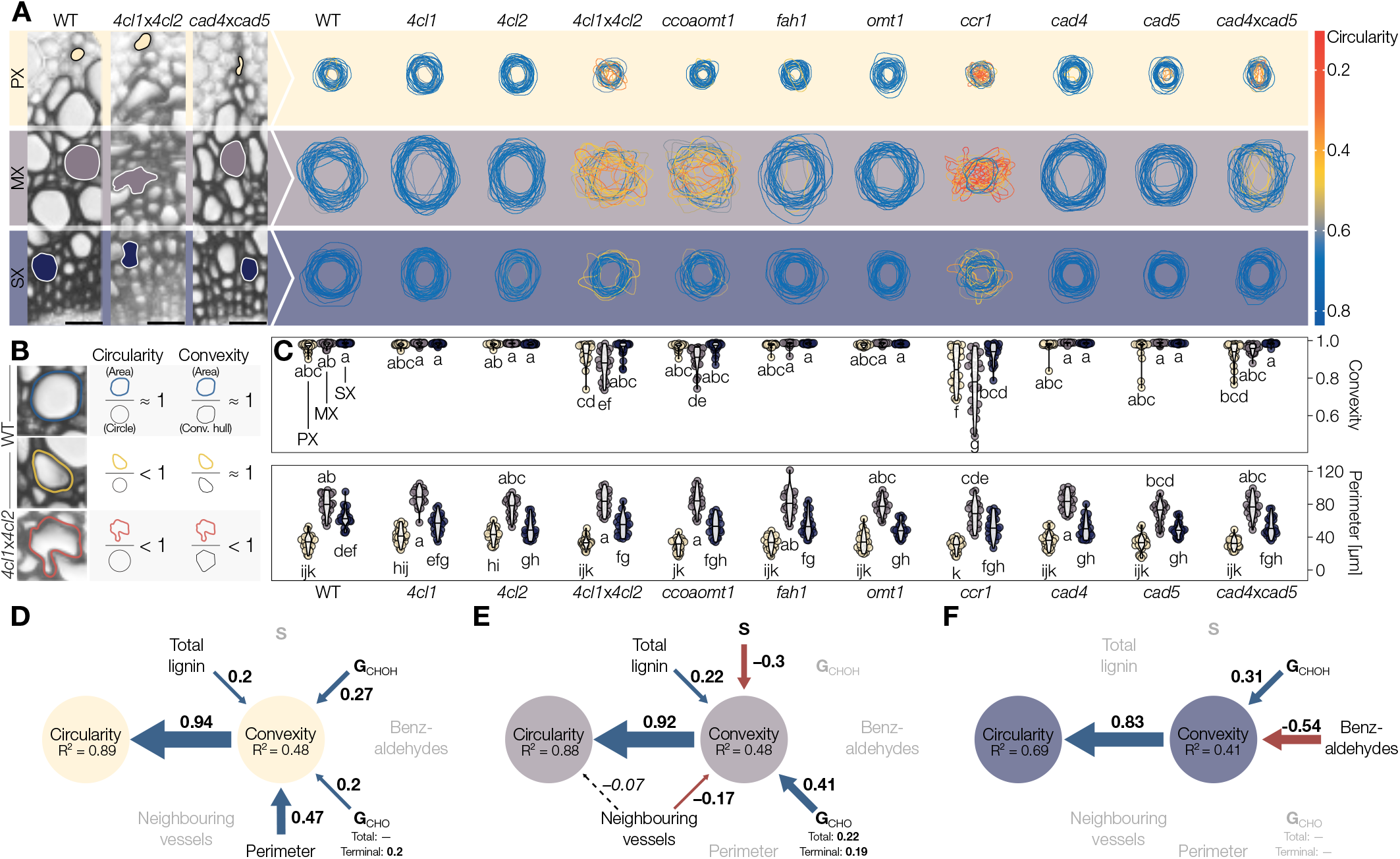
Lignin structure differently alters the resistance of specific TE morphotypes in annual plants. **A** Representation of 25 representative perimeters for each TE type in transverse cross-sections from stems of *Arabidopsis thaliana* loss-of-function mutants altered in lignin structure. TE outline colour indicates the circularity of each respective TE. **B** Schematic explanation of circularity and convexity of TEs. Any deviation from a perfect circle will decrease circularity, whereas only inward collapse of the perimeter will decrease convexity. **C** Convexity and perimeter of PX, MX and SX TEs in different phenylpropanoid biosynthesis mutants; *n* = 50 TEs per morphotype and genotype. Letters indicate significant differences according to a Tukey-HSD test (per panel; α = 0.05). **D–F** Structural equation models of the factors influencing TE convexity and circularity in the PX (**D**), MX (**E**) and SX (**F**) of *Arabidopsis thaliana*. Blue arrows and positive standardised coefficients indicate significant positive effects, red arrows and negative standardised coefficients indicate significant negative effects. Dashed arrows indicate predictors that were included and improved the model, but whose specific effects were not statistically significant. Greyed out variables did not improve the model and were excluded.

### *Post-mortem* incorporation of specific lignin residues alters TE resistance during wood formation in perennial plants

To assess the role of lignin during TE gradual *postmortem* maturation in tissues, we similarly evaluated TE lignification and resistance to collapse in woody whole plants of hybrid poplar. We focused our analyses on different TE types in stem cross-sections, primary (PV) and secondary xylem TEs/vessels (SV) at different developmental states, young and old SVs defined by the 50% distance to the cambium (Fig. 5A). All TEs in WT plants had similar surrounding cell types but varied in the lumen area, which was smallest in young SVs (Fig. 5B-C). *In situ* analysis of cell wall biochemistry showed differences in lignin levels (lowest in PVs) and **S**/**G** composition (highest in young SVs and gradually decreasing in old SVs and PVs) (Fig. 5D,E). Overall **G**_CHO_ levels were low, in accordance with the literature (Yamamoto et al. 2020), but the **G**_CHO_/**G**_CHOH_ ratio gradually increased from young SVs to PVs (Fig. 5F). Note that TEs differed between aspen and *Arabidopsis* presenting much larger perimeters for all morphotypes and different lignin composition, as previously shown (Chaffey et al. 2002; Agarwal et al. 2011), although still conserving the typical **G** residues enrichment (Figs. 5 and S6). This observation further confirms that lignin is adjusted for the specificities of each TE morphotype. Complete analysis of TE maturation across wood from the cambium to the pith showed gradual increases of lignin concentrations but also showed a reduction in **S**/**G** and an increase in **G**_CHO_/**G**_CHOH_ (Fig. S6). These results confirm that the process of TE *post-mortem* lignin accumulation observed in iPSCs also occurs in wood of whole plants, and show a dynamic change of lignin chemistry during these *post-mortem* processes as wood tissues mature.

**Figure 5.**
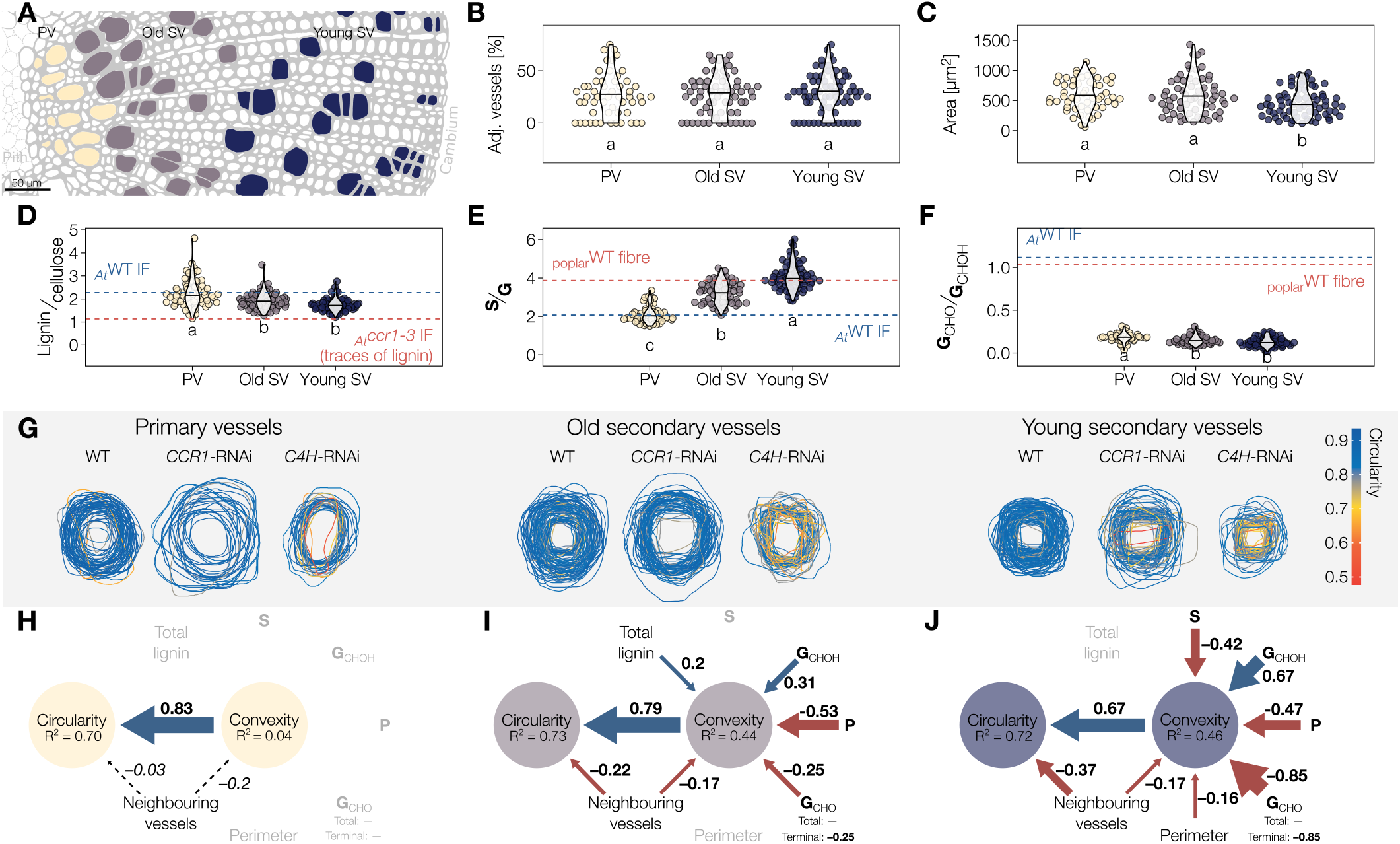
Different TE morphotypes in woody plants depend on specific *post-mortem* accumulated lignins for their resistance against collapse. **A** Scheme of the three TE types in the xylem of poplar stems, oriented on the pith–cambium axis: primary vessels (PV) in yellow, old secondary vessels (old SV) in purple and young secondary vessels (young SV) in blue. **B** Relative proportion of adjacent TEs for each TE type. Note that the proportion of TEs neighboring other TEs is independent of TE type and very similar to the proportions in *A. thaliana*. **C** Lumen area of each TE type determined from cross-sections. **D** Relative lignin to cellulose ratio measured by Raman microspectroscopy. **E** Relative **S** to **G** ratio measured by Raman microspectroscopy. **F** Relative **G**_CHO_ to **G**_CHOH_ ratio measured by Raman microspectroscopy. Because range and intercept of Raman band ratios differ from other biochemical analyses but still maintain a linear relationship (Agarwal 2019; Blaschek, Nuoendagula et al. 2020), references are presented for each lignin parameter using poplar fibres or the IF of *A. thaliana* WT and relevant mutants. Letters in panels **B–F** indicate significant differences according to a Tukey-HSD test (per panel; α = 0.05); *n* = 56–72 individual cells from 5 plants per TE type. **G** Representation of TE perimeter for each TE type in transverse cross-sections from stems of *Populus tremula*×*tremuloides* RNAi plants altering lignin biosynthesis. TE outline colour indicates the circularity of each respective TE. **H–J** Structural equation models of the factors influencing TE convexity and circularity in the PV (**H**), old SV (**I**) and young SV (**J**). Blue arrows and positive standardised coefficients indicate significant positive effects, red arrows and negative standardised coefficients indicate significant negative effects. Dashed arrows indicated predictors that were included and improved the model, but whose specific effects were not statistically significant. Greyed out variables did not improve the model and were excluded.

#### Lignin concentration and composition fine-tune the mechanical properties of TEs during their *post-mortem* maturation in woody tissue

To assess the link between lignin and TE collapse in wood, we followed the same strategy as in *Arabidopsis* using transgenic poplar with RNAi constructs to alter lignin amount and composition. TEs in the three genotypes used (WT, *CCR1*-RNAi, *C4H* -RNAi) also had a wide variation in lignin structure and collapse that moreover depended on their developmental age (Figs. 5G and S6). Using the single cell analyses of 100–170 individual TEs per morphotype and developmental stage, we could evaluate which parameters were associated with the capacity of specific TEs to withstand collapse using structural equation models. As PVs did not collapse, none of the measured parameters had any effect, thereby showing unique resilience of PVs (Fig. 5H). In old SVs resistance to collapse was associated with increases of **G**_CHOH_ and lignin levels but compromised by increases in terminal **G**_CHO_ residues and increases in neighbouring TEs (Fig. 5I). The resistance to collapse of young SVs was promoted by increases of **G**_CHOH_ but reduced by increases of **S**, terminal **G**_CHO_ residues, TE perimeter and neighbouring TEs (Fig. 5J). Similar to *Arabidopsis*, some of these effects were interdependent on each other. The effects of **G**_CHO_ residues in old SVs were affected by their adjacency to other vessels (Fig. S5), while in young SVs, the effects of **S** and **G**_CHO_ were similarly adjusted by each other and vessel perimeter/adjacency (Fig. S5). Altogether, these results confirm in woody tissues that the different lignin residues (**S**/**G** and **G**_CHO_/**G**_CHOH_), which are dynamically modulated during *post-mortem* maturation, have different impacts on the biomechanics of specific TE types to resist collapse.

### Aliphatic and aromatic changes in lignin residues nonredundantly control distinct mechanical properties of plant stems

To define how changes in lignin composition affected the biomechanics of TE cell walls and whole plants, we evaluated the impact of changes in the **G**_CHO_ to **G**_CHOH_ content of lignin. It showed opposite influence on TE collapse in both herbaceous and woody plants (Figs. 4,5) but also changed during TE *post-mortem* maturation (Fig. S6). We performed three point bending flexure measurements on *Arabidopsis* plant stems (Figs. 6 and S7; Nakata et al. 2020). We used three different segments along the stem length, from apices to bases, to spatially separate the lignification state of TEs and other vascular cells as previously performed (Fig. 6A; Pesquet et al. 2013; Hoffmann et al. 2020; Morel et al. 2022). We first evaluated the influence of turgor pressure and sap/water content on stem segments flexure after incubation in air, pure water or 1 M sorbitol for several hours. Reducing water content in stems did not alter stem flexural strength, flexibility or stiffness (Figs. 6C,D and S7), thereby showing that stem mechanical properties do not depend on water content but on cell walls. We then used two *A. thaliana* mutants altering lignin composition in **G**_CHO_ or **G**_CHOH_ to see how stem mechanical properties were affected. Mutants included *fah1*, devoid of **S** and instead accumulating **G**_CHOH_ (Meyer et al. 1998; Blaschek, Nuoendagula et al. 2020; Yamamoto et al. 2020), and *cad4*×*cad5* enriched in total and terminal **G**_CHO_ (Blaschek, Champagne et al. 2020; Sibout et al. 2005; Yamamoto et al. 2020). These biochemical changes differently affected TEs in whole stems of *fah1* and *cad4*×*cad5* compared to WT plants, lignin levels were slightly reduced but **S**/**G, G**_CHO_/**G**_CHOH_ and terminal/total **G**_CHO_ were largely altered (Fig. 6E). Similar biochemical changes in lignin were also be observed at the whole stem level (Fig. S7). Flexural strength and stiffness were significantly different between genotypes, with each mutant distinct from WTs, and affected more in apical than middle and basal stem segments (Fig. 6F-G).The changes included an increase of stiffness when increasing **G**_CHOH_ in mutant *fah1* and a decrease when increasing **G**_CHO_ in mutant *cad4*×*cad5* compared to WTs (Figs. 6F and S7). Similar reduction in stiffness due to increase of **G**_CHO_ in wood lignin had also been shown in poplar stem (Özparpucu et al. 2017; Özparpucu et al. 2018), revealing a conserved impact of **G**_CHO_ enrichment in lignin for herbaceous and woody species. In contrast, flexibility was unaltered between *fah1* and WT plants but increased by ∼3-fold in *cad4*×*cad5* (Fig. 6G). Our results therefore showed that the terminal aliphatic function of **G** residues in TEs had opposite effects on the cell wall biomechanics, increasing stiffness for alcohols compared to enhancing flexibility for aldehydes. Altogether, our results showed that the importance of regulating **G** residues with specific aliphatic terminal functions to differently modulate cell wall biomechanics, thus providing a mechanistic function behind the diversity of lignin residues.

**Figure 6.**
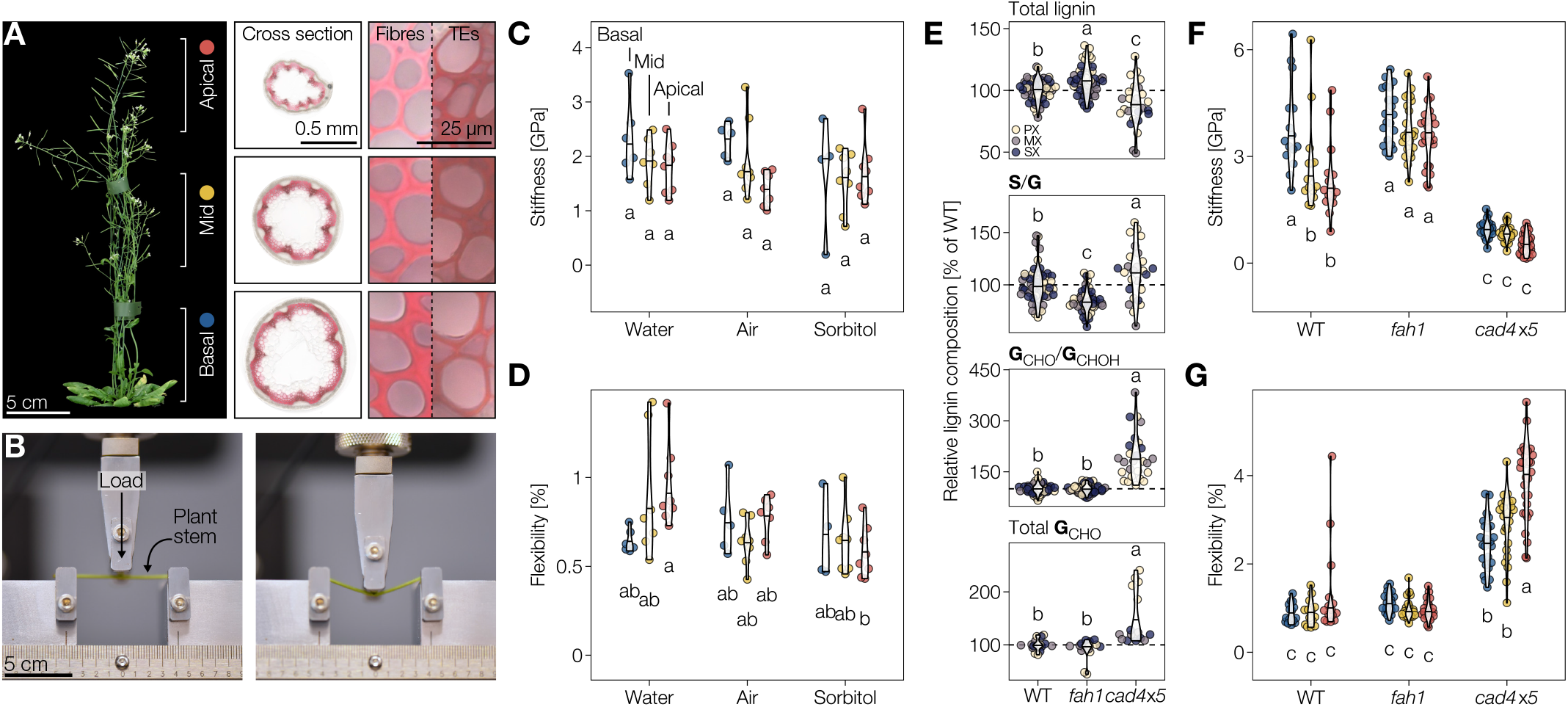
Distinct lignin monomers non-redundantly control specific mechanical properties. **A** Six-week-old *A. thaliana* WT plant with basal, middle and apical stem segments showing difference in TE developmental stages and marked with the colours representing them in subsequent panels. Wiesner stained cross-sections at the bottom of each segment with close-ups of interfascicular fibres and TEs shown. **B** *Arabidopsis thaliana* stem segment undergoing three-point-bending. Flexural behaviour is presented in movie S1. **C, D** The flexural stiffness (**C**) and sustained elastic deformation before irreversible breaking, *i*.*e*. flexibility, (**D**) of WT stem segments incubated in water, air, or sorbitol determined by three-point-bending; *n* = 4–8 stem segments per developmental stage and condition. **E** Total lignin, **S**/**G** and terminal **G**_**CHO**_/total **G**_**CHOH**_ (measured by Raman microspectroscopy) and total **G**_**CHO**_ (measured using the Wiesner test) in PX, MX and SX TEs of the different genotypes, expressed relative to the WT TEs of the respective morphotype; *n* = 15–50 TEs per genotype. **F, G** The flexural stiffness (**F**) and sustained elastic deformation before irreversible breaking (**G**) of stem segments from WT, **S** depleted *fah1* and **G**_**CHO**_ over-accumulating *cad4*×*cad5* mutant plants determined by three-point-bending; *n* = 14–30 stem segments per developmental stage and genotype. Letters in panels C–G indicate significant differences according to a Tukey-HSD test (per panel; α = 0.05).

### Increased TE cell wall flexibility due to coniferaldehyde in lignin enables plants to better recover from drought

As optimal vascular conduction is enabled by circular uncollapsed TEs (Zhang et al. 2016), we then evaluated how changes in biomechanics of TE cell walls affected their conductive function under drought conditions. More precisely, we estimated the effect of increased flexibility of TE cell walls, due to more **G**_CHO_ residues incorporation in lignin, would have on sap conduction capacity in response to normal watering and simulated drought using the osmoticum polyethylene glycol PEG_6000_ (Osmolovskaya et al. 2018). To ensure that the *cad4*×*cad5* mutation did not affect the lignin implicated for the endodermis function in water absorption/conduction, we monitored the apoplastic barrier of *cad4*×*cad5, fah1* and WT plants using propidium iodine on young seedlings (Y. Lee et al. 2013). No differences in apoplastic barriers were observed between the *cad4*×*cad5* and *fah1* mutants and WT plants (Fig. S8). Under normal conditions, 4–5-week-old WT plant rosettes had evapotranspiration rates of 7.5 mg water loss per min, which was significantly reduced by ∼25% under simulated drought when watered with 10% and 20% PEG_6000_ solution for 72h (Figs. 7A–C and S9). The lowered evapotranspiration confirmed the normal response of plants to drought, essentially due to stomatal closure (Martin-StPaul et al. 2017). Leaf wilting and chlorosis were visible under drought and accentuated by the level of PEG_6000_ treatment (Fig. 7A,B). Recovery experiments by transferring plants to normal watering for 96 h showed that plants in 20% PEG_6000_ did not recover whereas only 12.5% of plants in 10% PEG_6000_ fully recovered (Fig. 7D). Analysis of hypocotyl vasculature after re-watering showed a significant reduction in convexity increasing with PEG_6000_ levels (Fig. S9). This result indicated that drought in WT plants caused such strong inward collapse that TEs were unable to recover/restore their original shape after re-watering. In the *cad4*×*cad5* mutant, evapotranspiration rates also gradually decreased with increasing PEG_6000_ levels (Figs. 7A-C and S9). However, the mutant plants showed less wilting and chlorosis than the WT, resembling untreated plants (Fig. 7A-B). Recovery experiments to normal watering showed that 35-47% of *cad4*×*cad5* plants fully recovered after both 10% and 20% PEG_6000_ treatments (Fig. 7D). Analysis of hypocotyl vasculature in *cad4*×*cad5* plants after re-watering showed a slight by non-significant collapse of TEs with PEG_6000_ levels (Fig. S9). Overall, our results showed that the increased flex-ibility of TE cell walls, due to higher levels of **G**_CHO_ residues incorporation in lignin, either reduced TE capacity for irreversible inward collapse and/or increased TE capacity to recover initial shape, thus enabling plants to better resist drought. We therefore show that lignin composition in the TE cell walls directly influences their hydraulic properties and capacity to sustain and/or recover from extreme environmental changes.

**Figure 7.**
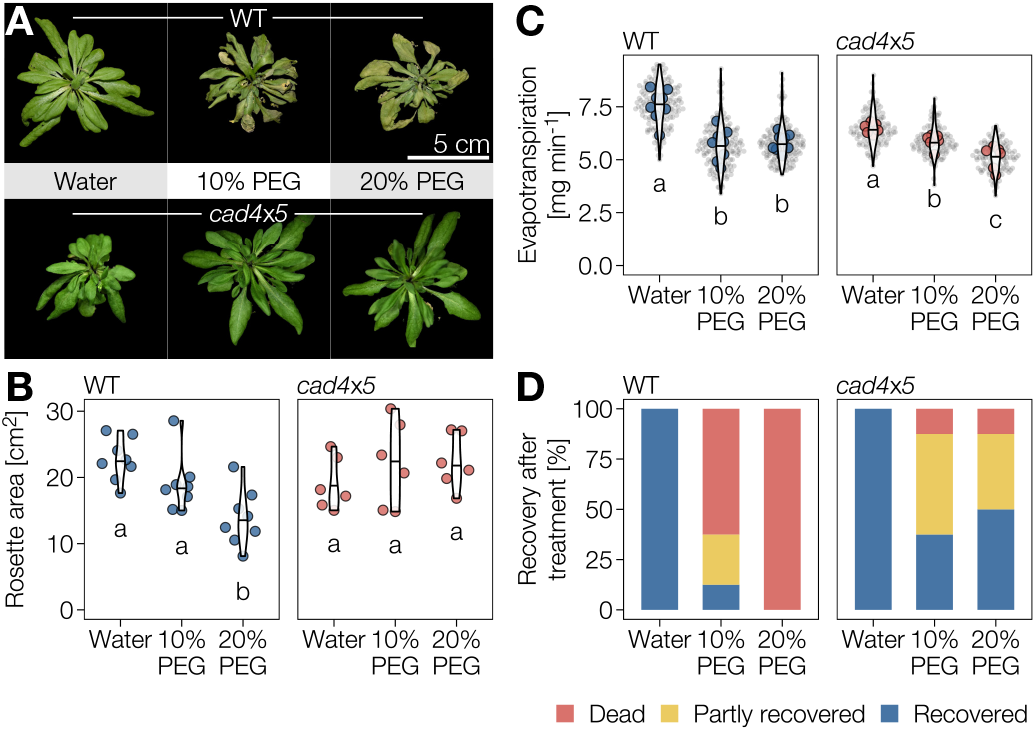
Coniferaldehyde induced flexibility of TE lignin improves plant resistance and/or recovery from extreme Ψ differentials. **A** Top view of 4-to 5-week-old *A. thaliana* WT and **G**_**CHO**_ over-accumulating *cad4*×*cad5* mutant plants after being irrigated with water, 10% PEG or 20% PEG for 3 days. **B** Top view rosette area after being irrigated with water, 10% PEG or 20% PEG for 3 days. Letters indicate significant differences according to a Tukey-HSD test (per panel; α = 0.05). **C** Evapo-transpiration rates of WT and *cad4*×*cad5* plants after being irrigated with water, 10% PEG or 20% PEG for 3 days. Small grey dots represent individual measurements, larger coloured dots represent the average per plant. Letters indicate significant differences according to a Tukey-HSD test (per panel; α = 0.05). **D** Proportion of WT and *cad4*×*cad5* plants that did not, partly or fully recover after being treated (with water, 10% PEG or 20% PEG for 3 days) by a 4 days recovery period in water-saturated soil.

### Increasing coniferaldehyde residues in lignin changes molecular conformations, torsions and reduces stiffness compared to coniferyl alcohol residues

To understand how **G**_CHO_ and **G**_CHOH_ residues differently contributed to lignin biomechanics in TEs at the molecular level, we performed molecular dynamic simulations of lignin oligomers differing in their **G** residues. As **G**_CHO_ and **G**_CHOH_ residues are mostly interlinked with β–*O*–4 ether linkages *in planta* (Yamamoto et al. 2020), we designed β–*O*–4 interlinked homomeric heptamers of **G**_CHO_ and **G**_CHOH_ (with or without αC–OH) according to previous characterisation studies (Grabber et al. 1998; Jourdes et al. 2007; Holmgren et al. 2009; Dima et al. 2015). Such lignin oligomers composed of only **G**_CHO_ or **G**_CHOH_ have been chemically synthesised *in vitro* (Önnerud et al. 2002; Ito et al. 2002; Holmgren et al. 2009; Notley and Norgren 2012; Koshiba et al. 2013; Blaschek, Nuoendagula et al. 2020). Molecular dynamic simulation showed that oligomers of **G**_CHO_, which maintain an unsaturation in β–*O*–4 linkages (Grabber et al. 1998; Jourdes et al. 2007; Holmgren et al. 2009), greatly reduced rotation of the _α_C–_β_C torsions but not the rotation around _β_C–O or C_6_–O–_α_C compared to **G**_CHOH_ with or without αC–OH (Fig. S10A,B). Differences in molecular volume and density showed slightly larger and denser polymers for **G**_CHOH_, independently of αC–OH, compared to **G**_CHO_ (Tab. 1). The conformation of **G**_CHOH_ oligomers, independently of the αC–OH, were however substantially more compact with significantly smaller radius of end-to-end distance (Re-e) and of gyration (Rg) than the more extended conformations for **G**_CHO_ oligomers (Tab. 1). Thesefindings showed the important influence of different terminal functions in the aliphatic chain of lignin residues, leading to polymers with very different conformations and thus affecting their capacity to fold and pack in the available space between the polysaccharides of plant cell walls during *post-mortem* lignification (Fig. S10C–E). Our results showed that increases in **G**_CHO_ reduced the capacity of lignin polymers to compactly fold in these free cell wall spaces compared to **G**_CHOH_. We then performed molecular dynamic simulation of these different polymers under stress (external pressure) to evaluate the cumulative mechanical properties of 100 lignin oligomers made either of **G**_CHO_ or **G**_CHOH_ residues, with or without αC–OH. Significant increases in Young modulus (determining the relative stiffness to elasticity of any material) were observed, independently of αC–OH, for **G**_CHOH_ compared to **G**_CHO_ (Tab. 1). These results showed that the terminal function of lignin residues directly affected the biomechanics of each polymer and confirmed the differences in stiffness and flexibility observed in TEs and whole plants when modulating lignin chemistry (Figs. 4–6). Our molecular dynamic simulation analyses confirmed that the modulation of **G** residue terminal aliphatic functions directly controls lignin biomechanics.

**Table 1.** Properties of lignin heptamers computed in molecular dynamics simulations. Re-e is the root mean square of the end-to-end distance and σ_Re-e_ is its variance; Rg is average radius of gyration and σ_Rg_ is its variance.

| Oligomer | Volume [$\text{\AA}^3$] | Density [$\text{g cm}^{-3}$] | Re-e [$\text{\AA}$] | $\sigma_{\text{Re-e}}$ [$\text{\AA}$] | Rg [$\text{\AA}$] | $\sigma_{\text{Rg}}$ [$\text{\AA}$] | Young modulus [MPa] |
| --- | --- | --- | --- | --- | --- | --- | --- |
| $\text{G}_{\text{CHO}}$ | 169 | 1.21 | 19.6 | 8.5 | 9.5 | 1.7 | 1400 ± 200 |
| $\text{G}_{\text{CHOH}}$ | 174 | 1.205 | 13.6 | 5.9 | 7.25 | 0.75 | 2000 ± 200 |
| $\text{G}_{\text{CHOH}}$ with $\alpha\text{C-OH}$ | 179 | 1.25 | 14.1 | 5.4 | 7.8 | 1.2 | 1900 ± 200 |

## Discussion

TEs belong to the few cell types that fulfill their function only after their death, which greatly limits their capacity to adapt to changing environmental or developmental constraints. Accordingly previous studies assumed that functional TEs were inert and had no adaptive capacity. We however showed that TEs can still adapt after death by accumulating lignin *post-mortem* in their SCWs (Figs. 1, 5 and S6). This process occurred in TEs of both annual and perennial species (Figs. 1, 5 and S6), thereby setting *post-mortem* lignification as a conserved mechanism of TEs made to reinforce their conducting roles once hollowed by cell death in many vascular plants. We moreover showed that TE *post-mortem* lignification not only increased total lignin concentration but also changed composition as TE matured (Fig. S6). We further confirmed that TE cell walls are specifically enriched in **G** residues alike other tracheophytes (Pesquet et al. 2019), and showed that each TE morphotype has a distinct lignin composition which cannot be limited to **S**/**G** ratio (Figs. 3 and 5). This specific cell wall lignin composition for each TE morphotype was essential to prevent inward collapse for an optimal hydraulic conductivity, as increased accumulation of **S** instead of **G** residues comprised the biomechanics of most morphotypes. This result complemented recent observations that showed collapse of MX in *Arabidopsis* when genetically increasing their **S** content (Sakamoto et al. 2020). This morphotype specific resolution of lignin highlights the benefits of using *in situ* methods capable of measuring biochemical and biomechanical aspects at the single cell and even cell wall levels, such as SEM-EDS, AFM as well as Raman spectroscopy (Figs. 1–6). Unlike biochemical methods using ball-milling and grinding, these *in situ* quantitative imaging methods avoid averaging errors and compound effects between cell types at different maturation stages when analysing whole plant organs. However, the direct analysis of lignin *in situ* using Raman spectroscopy does not enable to reliably determine **H** residue proportion or the different aliphatic functions of **S** residues (Blaschek, Nuoendagula et al. 2020). We show that the different aliphatic terminal function of **G** residues, such as alcohol or aldehyde, are specifically accumulated in different concentrations between TE morphotypes (Figs. 4,5) during TE *post-mortem* maturation, with a delayed deposition of **G**_CHO_ compared to **G**_CHOH_ (Fig. S6; Blaschek, Champagne et al. 2020; Kutscha and Gray 1972). Specific control of **G** _CHOH_ and **G** _CHO_ accumulation in lignin had previously been shown between wood cell types and cell wall layers in multiple plant species (Peng and Westermark 1997; J. S. Kim et al. 2015; Zheng et al. 2016; Blaschek, Champagne et al. 2020) and in response to environmental conditions (Euring et al. 2012; Camargo et al. 2014; Nakagawa et al. 2012). We showed that each TE morphotype requires a specific **G** _CHO_/**G**_CHOH_ compositional ratio to fine-tune the stiffness to flexibility of their cell walls in single cells as well as in whole stems (Figs. 2–6 and S7). The adjustment of stiffness to flexibility of ether-linked **G** residues only depended on the terminal aliphatic function as shown by our molecular dynamic simulations (Tab. 1; Fig. S10). As TE dimensions and cellular surrounding greatly vary between plant species, our results moreover show that each plant species fine-tunes the **G** _CHO_/**G**_CHOH_ with other changes in lignin chemistry for their distinct TEs (Figs. 3–5, S4–6). In addition to the role of lignin chemistry to support TE function, our results confirmed that the vascular tissue organisation (TEs neighbouring other TEs) affects TE biomechanics and hydraulics but only for specific morphotypes (Figs. 4-5; Lens et al. 2011; Martínez-Vilalta et al. 2012). Changes in cell wall stiffness to flexibility at the TE level might represent a mechanism to adapt TE biomechanics to environments where water availability varies brutally, thus enabling TEs to regain their original shape after collapse to restore plant sap conduction. We propose that the trade-off between stiffness and flexibility of TEs is regulated by controlling lignin composition. Crucially, this regulation continues after TE cell death, showing that TEs are not an inert arrangement of pipes, but a dynamic system that can continuously adjust its properties through *post-mortem* lignification. The exact identity and function of the different components enabling TE *post-mortem* lignification such as the living cooperating cell types, the metabolites used and molecular actors regulating lignin formation through biosynthesis, transport and/or polymerisation still need to be identified. Our study therefore establishes that the proportions of the different lignin residues, varying in both aromatic and aliphatic substitutions, are specifically regulated in each cell type and along their maturation to enable specific cellular functions.

## Materials & methods

### Inducible pluripotent cell suspension cultures (iPSCs)

*Arabidopsis thaliana* iPSCs were produced and induced to differentiate into isolated tracheary elements as previously described (Pesquet et al. 2010; Ménard et al. 2017). Cell suspensions were induced by adding hormones to 30 mg ml-1 of 10 days old cells (fresh weight) in 1x Murashige and Skoog (MS) medium (Duchefa, M0222.0025) at pH 6.0 with 10 μM of morpholino-ethanesulfonate (Sigma-Aldrich, M8250) and 3% sucrose. Xylogenic induction was triggered by adding 3.2 nM α-naphthaleneacetic acid (Sigma Aldrich, N0640), 44.4 nM 6-benzyl-aminopurine (Sigma Aldrich, B3408) and 4 μM 24-epibrassinolide (Sigma Aldrich, E1641). The inhibition of lignin monomer biosynthesis was performed by adding 12.5μM of piperonylic acid (Sigma-Aldrich, P49805) at the time of hormonal induction of TE differentiation in suspension cultures as previously described (Van de Wouwer et al. 2016; Decou et al. 2017).

### Plant material

*Arabidopsis thaliana* and hybrid poplar plants were grown in climacteric growth chambers under a 16 /8 h long day light regime with 150 μmol m-2 s-1 illumination using (Aura T5 Eco Saver Long Life HO light tubes; Aura-Light, Sweden) and 22°C/18°C in 60% humidity. *Arabidopsis thaliana* mutants in the Columbia *Col-0* background used included *ccoaomt1* (SALK_151507; Kai et al. 2008), *fah1* (EMS mutant; Meyer et al. 1998), *omt1* (SALK_135290; Tohge et al. 2007), *4cl1-1* (SALK_142526; Van Acker et al. 2013), *4cl2-4* (SALK_110197; Li et al. 2015), *4cl1*×*4cl2* (Blaschek, Champagne et al. 2020), *ccr1-3* (SALK_123-689; Mir Derikvand et al. 2008), *cad4* (SAIL_1265_A06; S. Lee et al. 2017), *cad5* (SAIL_776_B06;S. Lee et al. 2017) and *cad4*×*cad5* (Blaschek, Champagne et al. 2020). *Populus tremula*×*tremuloides* hybrid poplars clone T89 were transformed as described by Nilsson et al. (1992) with RNA interference constructs targeting either *CINNAMATE-4-HYDROXYLASE* (Potri.013G157900; Bjurhager et al. 2010) or *CINNAMOYL-COA REDUCTASE* (Potri.003G181400) selected for best reduced gene expression (Escamez et al. 2017). Poplar plants were soil-grown in conditions identical to those described above.

### Atomic Force Microscopy (AFM)

AFM imaging was performed on cell samples semi-dried for less than 1 hour using a Dimension Icon AFM (Bruker, Nanoscope controller, Santa Barbara, CA, USA). The measurement was conducted under Peak-Force QNM mode in air condition by using the probe TESPA-V2 (Bruker). The force set-point was 0.15V. The height, peak-force error, DMT (Derjaguin-Muller-Toporov) modulus, adhesion, and deformation images were recorded after calibrating the probes on Mica. The images were processed by NanoScope Analysis 1.5 software (Bruker) and quantification performed using ImageJ distribution Fiji (Schindelin et al. 2012).

### Scanning electron microscopy (SEM) coupled with Energy-dispersive X-ray spectroscopy (EDS)

Cell samples for electron imaging and chemicals analysis were dispersed and sedimented on glass coverslips, then either (i) dehydrated in a series of graded ethanol and critical point dried (CPD) using a Leica EM CPD300 critical point dryer or (ii) submitted to air drying, and finally coated with 5 nm chromium using Quorum Technologies Q150T ES metal coater. The samples morphology was analyzed by field-emission scanning electron microscopy (SEM; Carl Zeiss Merlin) using an in-lens secondary electron detector at accelerating voltage of 4 kV and probe current of 100 pA. Elemental composition measurements were performed using an energy-dispersive X-ray spectrometer (EDS; Oxford Instruments X-Max 80 mm2) at accelerating voltage of 10 kV and probe current of 300 pA, where the elemental composition percentage is an average of multiple line and point analyses.

### Histological preparation and analyses

Eight-week-old in-florescence stem bases or four–five-week-old hypocotyls were cleared in 70% ethanol, rinsed in water and embedded in 10% agarose prior to sectioning to 50 μm with a VT1000S vibratome (Leica, Sweden). TE cell wall autofluorescence was acquired using LSM800 (Zeiss, Germnay) confocal microscope with excitation at 405 nm and emission collected from 420-650 nm (Decou et al. 2017). Semi-quantitative Raman microspectroscopy was performed as described by Blaschek, Nuoendagula et al. (2020) on the different vessel types using a confocal Raman microscope (RAMANplus, Nanophoton, Japan and LabRAM HR 800, Horiba, France) with a 532 nm laser. Averaged spectra were obtained from three to seven cell walls per TE morphotype and plant, in one to three plants per genotype for *Arabidopsis* and from 17 to 71 cell walls per TE morphotype and plant, in two to six plants for poplar. Quantitative Wiesner test was performed as described by Blaschek, Champagne et al. (2020) using an Olympus BX60 brightfield microscope equipped with an Olympus UPFLN 40X objective (NA 0.75), an Olympus XC30 CCD colour camera. TE morphological features (distance from cambium, lumen area, perimeter, circularity, neighbouring cell types) were measured from microscopy images using the ImageJ distribution Fiji (Schindelin et al. 2012). TE circularity was determined as 4π (area / perimeter2), and TE convexity as area / area of convex hull. Fiji macros are available at https://github.com/leonardblaschek/fiji.

### Three point flexural test

The stiffness and strength of stems were assessed using three-point flexural tests with an Instron 5966 universal testing machine (Instron, USA) equipped with a 100 N load cell in a humidity and temperature controlled room (50% RH and 23°C). 4–5 cm long stem segments from 25–35 cm stems of 6–7 week-old plants were placed on two supporting pins that were separated at an average span-to-diameter ratio of 38–39±4. Treatment to alter stem water content included incubation for several hours prior to bending in pure distilled water or 1M sorbitol (Sigma, S1876) solutions, all other measurements were performed in air. After manually lowering the loading pin until just in contact with the sample, the probe was lowered automatically at a constant displacement rate of 2 mm min-1 until a final displacement of 7 mm (movie S1). Th eflexural strength *σ*_max_ (MPa) was calculated as the maximumflexural stress using Eq. 1.

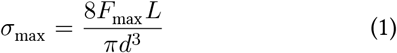

In this equation, *F* _max_ (N) is the maximum force the specimen can withstand before kinking, *L* (mm) is the span length between the supporting pins, and *d* (mm) is the diameter of the circular cross-section of the specimen that was determined using optical microscopy imaging. The flexural stiffness *E* (MPa) was calculated from the slope of the initial linear part of theflexural stress-strain curve using Eq. 2.

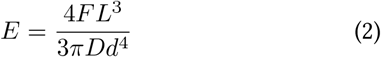

In this equation, *D* (mm) is the maximum deflection of the centre of the stem. The flexibility of stems was defined as the strain at maximum stress, *i*.*e*. the amount of deformation a stem could endure before irreversibly breaking.

### Evapotranspiration and simulated drought

Simulated drought treatments were made by watering plants with 0, 10 or 20% polyethylene glycol (PEG) 6000 (Sigma-Aldrich, 8.07491) in tap water for 72h in growing conditions (150μE light, 25C, 60% relative humidity). Evapotranspiration was measured using mass difference for 20 minutes on a LA-124i microbalance (VWR) directly connected to a computer and monitored using the i-Weight software (VWR). Plant recovery was done by placing potted plants directly in water for 72h. Light intensity, temperature and relative humidity were constantly monitored during the course of the measurements (Fig. S9). Images of the rosettes were acquired with Nikon D750 camera equipped with a 50mm F1.4 DG HSM lens. Image segmentation and rosette area measurements were performed in Fiji (Schindelin et al. 2012). Hypocotyl vessel collapse in normal and simulated drought was estimated as described above.

### Apoplastic connectivity staining

*Arabidopsis thaliana* seeds were surface sterilised (2 min 70% ethanol followed by 5 min 5% bleach) and stratified in water for 2 d at 4°C. Seeds were plated on 0.5x MS pH 5.7 with 0.8% agar and put in a growth chamber at 16/8 h day/night regime with 150 μmol m-2 s-1 illumination (Aura T5 Eco Saver Long Life HO light tubes; AuraLight, Sweden) and 22°C/18°C in 60% humidity. Seedlings were grown vertically until 4 days past germination. Formation of the functional apoplastic barrier was analysed using propidium iodide (PI; Sigma-Aldrich P4170) as described previously (Y. Lee et al. 2013). Briefly, seedlings were stained in 15 μM PI in deionised water for 10 min, rinsed twice in tap water, mounted in tap water between glass slide and cover slip and imaged using a Zeiss LSM 780 confocal microscope equipped with a 20x objective. The PI staining the apoplastic space was excited with a 488 nm laser and visualised by long pass emission at >500 nm. Tiles were stitched and analysed in Fiji (Preibisch et al. 2009; Schindelin et al. 2012). For quantification, cells were counted from onset of elongation (defined as cells being more than twice as long as they are wide) to the absence of any stain from the vascular cylinder.

### Lignin biochemical analysis

Lignin concentration in cell wall was determined after cell wall isolation according to Yamamoto et al. (2020) and the thioglycolic acid lignin derivatisation as described by Suzuki et al. (2009) on isolated extractive-free cell walls. Absorbance was measured at 280 nm and calibrated using a regression curve obtained using different quantities of alkaline spruce lignin (Sigma Aldrich, 471003). Pyrolysis-GC/MS was used to measure **S**/**G** and **G**_CHO_/**G**_CHOH_ according to Gerber et al. (2012) on 60 μg (±10 μg) of 8 week-old stem samples. Thioacidolysis-GC/MS-FID was used to determine the terminal/total positional ratio of β-*O-*4 linked **G**_CHO_ residues on 5 mg (±1 mg) of isolated cell wall from 8-week-old stem samples as described by Yamamoto et al. (2020).

### Data analyses and structural equation modelling

Data analysis and visualisation were performed in R (v4.0.4), using the “tidyverse” collection of packages (v1.3.0). The structural equation models were built using the “piecewiseSEM” package (v2.1.0; Lefcheck (2016)). The included variables were measured for each individual TE, except for the Wiesner test intensity in *A. thaliana*, for which the average per individual plant and TE type was used. The multiple linear regression models in the structural equation models were selected using a bidirectional step-wise optimisation approach, excluding interaction terms for clarity. Significantly contributing interaction terms were identified separately and visualised using the “interactions” R package (v1.1.1). Model fits and coefficients are summarised in table S3 and S4. R code used in this study is available at https://github.com/leonardblaschek/Rscripts/blob/master/irx_pub_figs.rmd.

### Molecular dynamic simulations

Molecular dynamics simulations were carried out on lignin heptamers made of either only **G**_CHOH_, with or without αC–OH, or **G**_CHO_ interlinked by β–*O*–4 ether linkages according to previous analyses (Grabber et al. 1998; Holmgren et al. 2009; Yamamoto et al. 2020). Molecular structures were designed with the Avogadro software (Hanwell et al. 2012) and processed using ACPYPE utility (Sousa da Silva and Vranken 2012) to produce GROMACS (Abraham et al. 2015) topology files implementing Generalised Amber Force Field (GAFF) (Wang et al. 2004). All simulations have been carried out using GROMACS v.2020 software. First, 100 molecules of the selected type were placed randomly in a cubic box of size 20 nm in random orientations. Short energy minimisation run was carried out to remove possible molecule overlap. This was followed by 1 ns run at 100 bar pressure to push the oligomers close to each other, then 400 ns run at 1 bar pressure with isotropic Berendsen barostat (Berendsen et al. 1984), and then 400 ns with anisotropic barostat. The obtained system was considered as equilibrium and used for further analysis. Young modulus computations were carried out by applying pressure between 100 and -200 bar with step 50 bar, in each of 3 directions. At each pressure step, a 300 ns simulation was carried out, the last 100ns were used to determine average box extension. The Young modulus was determined from the slope of strain-stress plot averaged in each of the 3 spatial directions. Simulations were repeated by starting from another random configuration and using the same protocol. Other relevant simulation parameters included: temperature control with Berendsen thermostat at T = 298°K and relaxation time 1 ps, time step 2 fs, constraint bonds to hydrogen atoms, Particle mesh Ewald summation of electrostatic interaction (Darden et al. 1993).

## Acknowledgements

We thank Anaxi Houabert, Louis Leboa, Charilaos Dimotakis and Drs, Elena Subbotina, Masanobu Yamamoto and Junko Takahashi-Schmidt for growing plants and providing lignin analysis in the different mutants. We thank Dr. Stefano Manzoni, Prof. Katharina Pawlowski and Prof. Hervé Cochard for advice and critical comments. We also thank the National Institute for Materials Science (NIMS) for access to Raman confocal microscope, the Albanova NanoLab for access to the AFM, the National Microscopy Infrastructure (NMI) for access to SEMEDS. This work was supported by Gunnar Öquist fellowship from the Kempe foundation (to EP), Vetenskapsrådet (VR) research grants 2010-4620 and 2016-04727 (to EP), the Stiftelsen för Strategisk Forskning ValueTree (to EP), the Bolin Centre for Climate Research RA3, RA4 and RA5 “seed money” and “Engineering Mechanics for Climate Research” (to EP), and the Carl Trygger Foundation CTS 16:362/17:16/18:306/21:1201 (to EP). Raman microspectroscopic analysis on plant cross sections was supported by the NIMS Molecule & Material Synthesis Platform in the “Nanotechnology Platform Project” operated by the Ministry of Education, Culture, Sports, Science and Technology (MEXT), Japan. We also thank Bio4Energy (a strategic research environment appointed by the Swedish government), the UPSC Berzelii Centre for Forest Biotechnology, the Institute of Global Innovation Research (GIR) of Tokyo University of Agriculture and Technology (TUAT), and the Departments of Organic Chemistry, of Materials and Environmental Chemistry (MMK), of Ecology, Environment and Plant Sciences (DEEP) and the Bolin Centre for Climate Research of Stockholm University (SU).

## Author contributions

EP conceived the study. EP and DM designed the experiments. DM, LBl, KK, CZ, HS, N, CCL, AL, ZB and EP performed the experiments. DM, LBl, KK, CCL, AL and EP analysed the data. LBe, AM, ZB, SK and EP ensured financial support and scientific expertise. EP wrote the article. All co-authors revised the manuscript.

