## Supplementary material for "Plant biomechanics and resilience to environmental changes are controlled by specific lignin chemistries in each vascular cell type and morphotype"

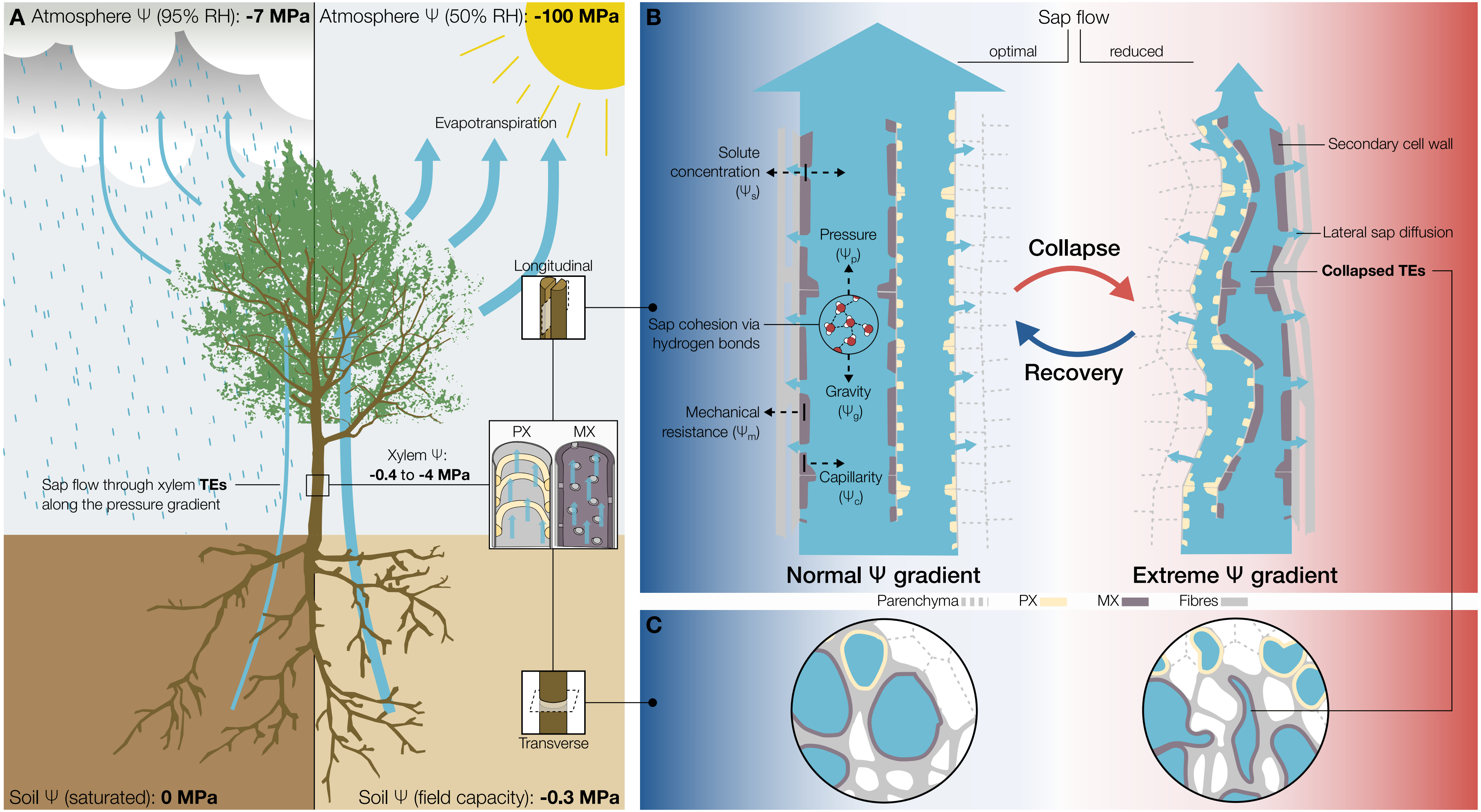


**Figure S1 | Introduction to basic physiological concepts of water conduction in TEs. A** The gradient of $\Psi$ between soil, plant and atmosphere that drives water transport through TEs depends on environmental conditions. **B, C** Schematic longitudinal (**B**) and transverse (**C**) sections through the different TE morphotypes. The total $\Psi$ is the sum of physical ($\Psi_c$, $\Psi_g$, $\Psi_m$, $\Psi_p$) and chemical ($\Psi_s$) pressures. Under normal conditions, TEs can withstand $\Psi$ without inward collapse, but extreme environmental changes and the associated large gradient in $\Psi$ leads TEs to collapse which affects water flow. Once the conditions return to normal, collapsed TEs can regain their shape.

MX, metaxylem; PX, protoxylem; RH, relative humidity; TE, tracheary element.

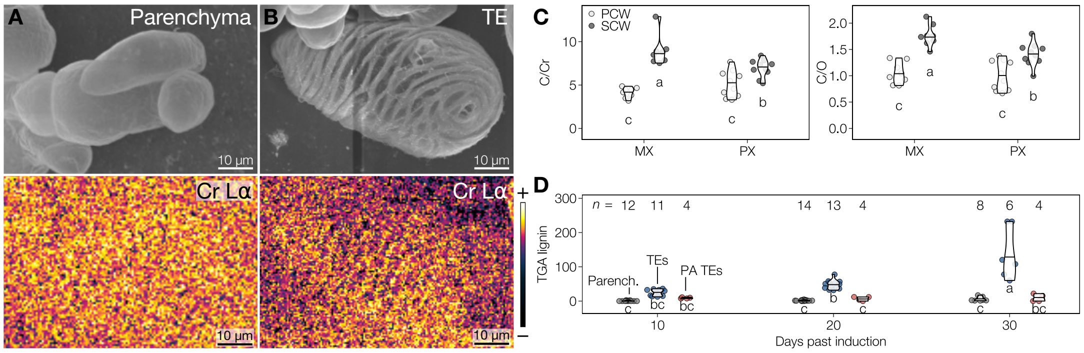


**Figure S2 | MX and PX TEs continuously lignify *post-mortem* (related to figure 1).** **A** Scanning electron micrograph of isolated parenchyma cell in uninduced condition, prepared using critical point drying (CPD), as well as its energy-dispersive X-ray spectroscopy (EDS) chromium (Cr) signal in color-coded intensity. **B** Scanning electron micrograph of isolated TE after 30 days in induced condition, prepared using CPD, as well as its EDS Cr signal in color-coded intensity. **C** C/CR and C/O ratios in PX and MX TEs from the plateau phase of cell wall lignification (>30 d after induction); $n$ = 8–10 individual cells per morphotype. **D** Lignin content of extracted cell walls from parenchyma cells, lignifying TEs and TEs treated with piperonylic acid (PA, 12.5 µM) as determined by thioglycolic acid derivatisation (TGA); $n$ = 4–14 independent cultures per time point and treatment.
Letters indicate significant differences according to a Tukey-HSD test (per panel; $\alpha$ = 0.05).

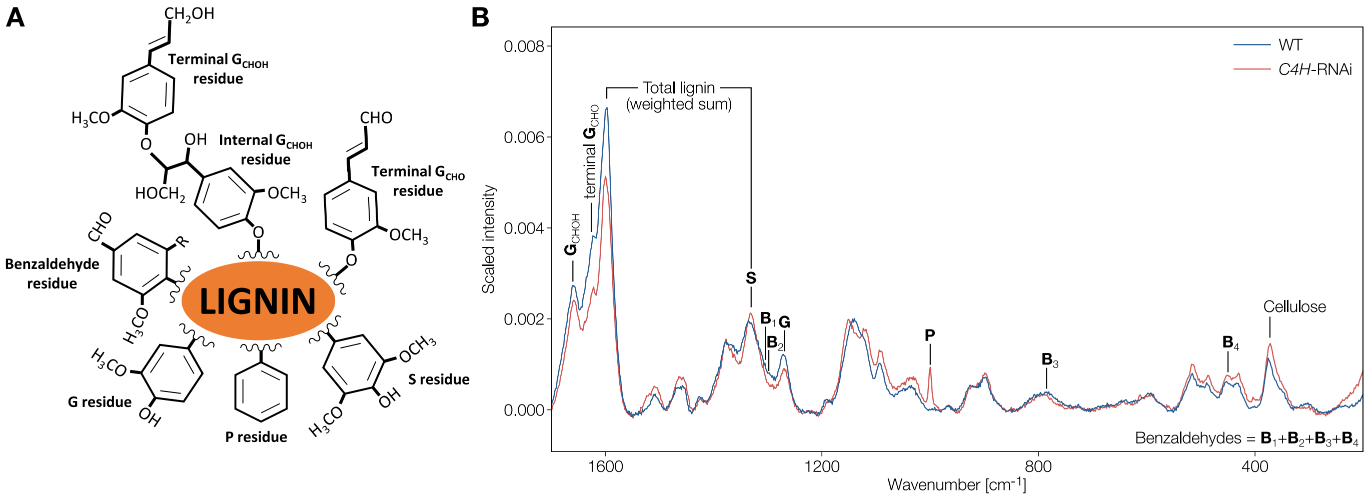


**Figure S3 | *In situ* quantification of lignin chemistry by Raman microspectroscopy (related to figures 3–6).** **A** Lignin residues detected by Raman microspectroscopy. **B** Raman bands used for the quantification of different cell wall polymers and lignin residues, shown on the example of representative spectra of TEs from *Populus tremula*×*tremuloides* WT and *C4H*-RNAi plants. The scattering of **P** (styrene) residues at 1000 $cm^{-1}$ is described in Noda and Sala (2000).

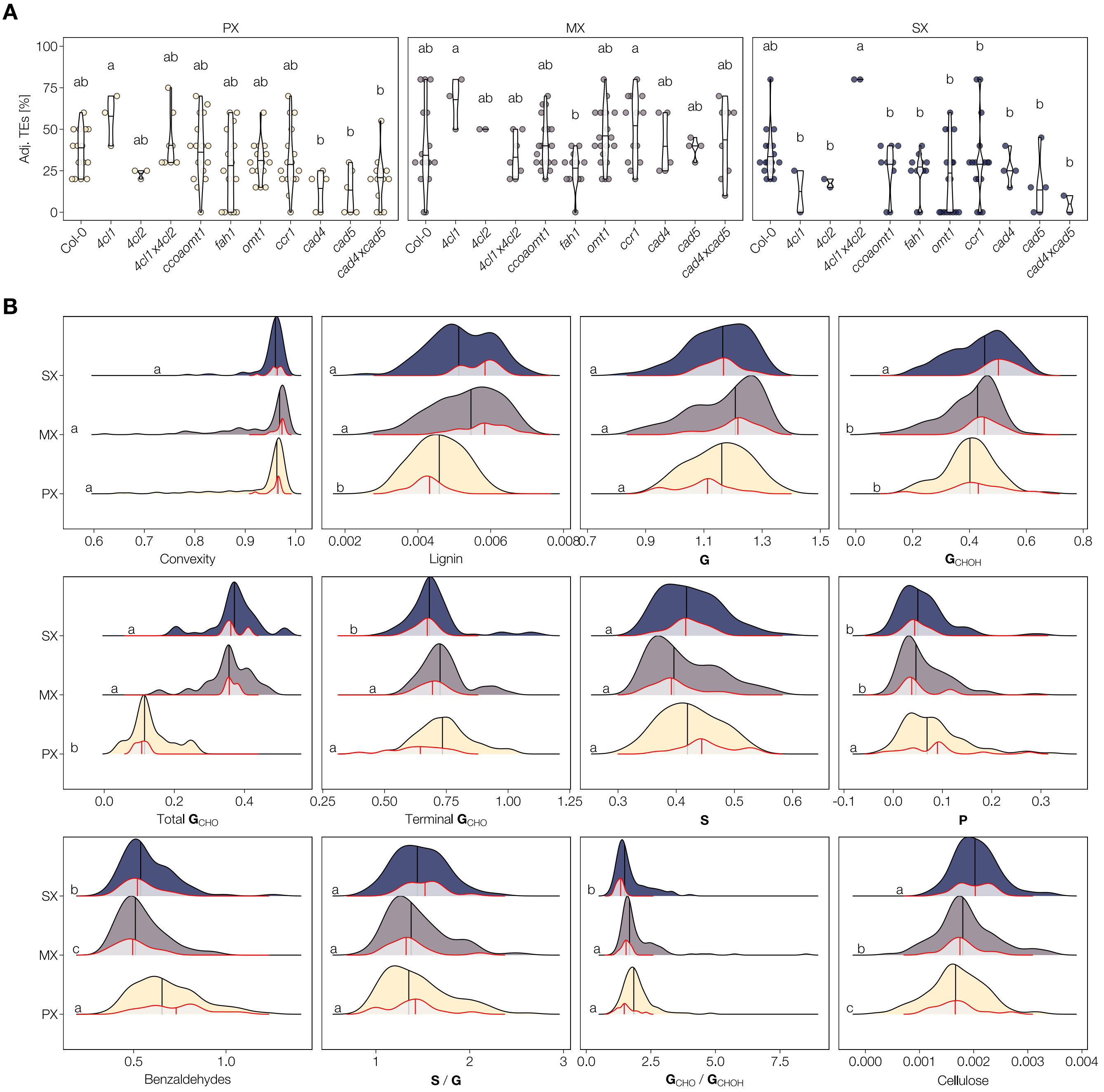


**Figure S4 | TE morphology and lignin composition in *A. thaliana* phenylpropanoid mutants (related to figure 4). A** Relative proportion of cell types surrounding TE perimeters other than TEs in the different *A. thaliana* genotypes comprising our dataset. Letters indicate significant differences between genotypes according to a Tukey-HSD test (per panel; $\alpha$ = 0.05). **B** *A. thaliana* TE convexity and cell wall composition data used in the structural equation models (Fig. 4 B–D). Variation across all genotypes (in blue/purple/yellow) overlayed with the variation in the WT (grey with red outline, scaled to 30%). Vertical lines represent the respective median values. Letters indicate significant differences between TE morphotypes according to a Kruskal-Wallis test followed by Dunn's multiple comparison (per panel; $\alpha$ = 0.05). All data used in the models is also available in the supplemental data file.

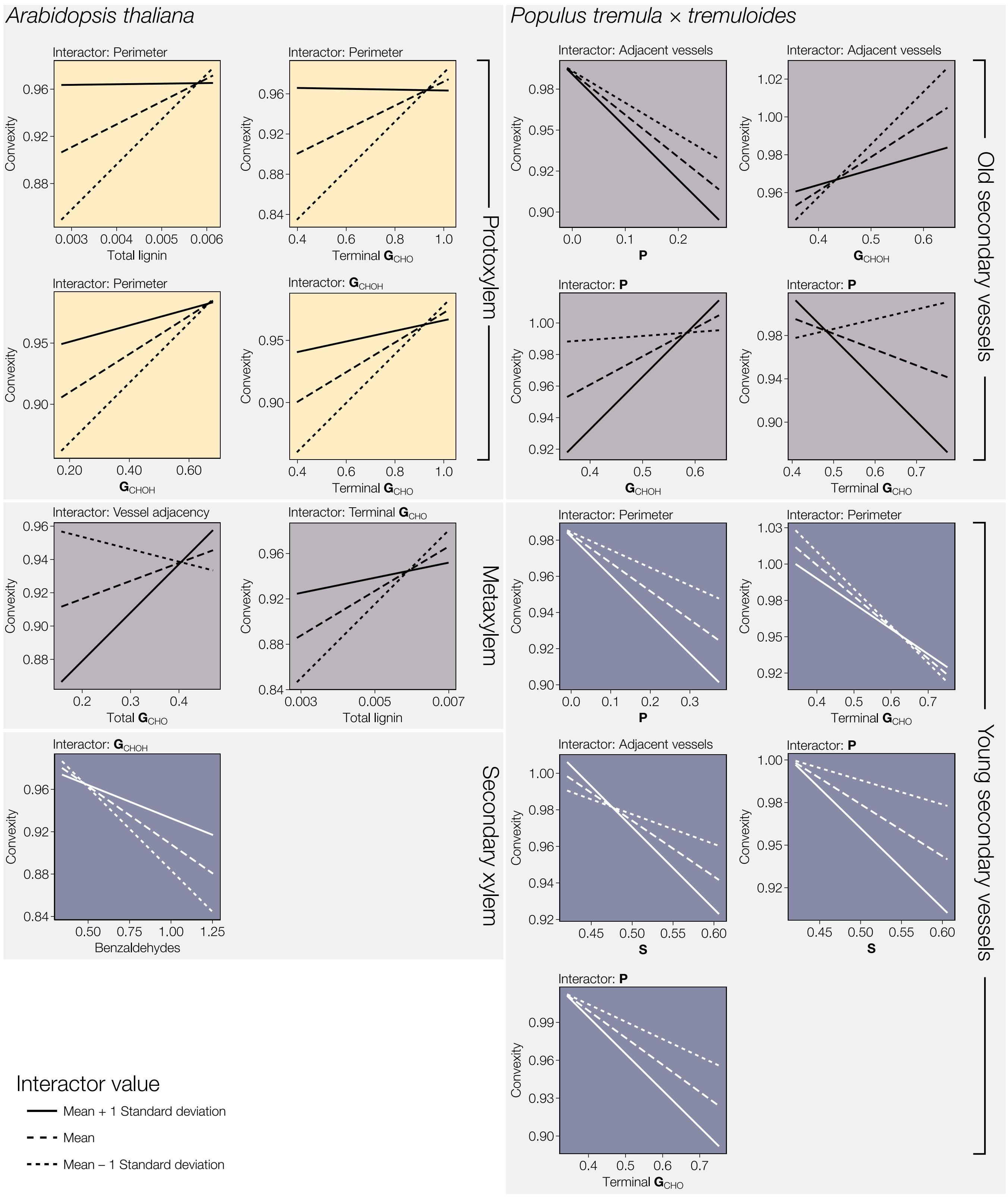


**Figure S5 | Effects of cell wall morphology and composition on convexity are interdependent (related to figures 4 and 5).** Several two-way interactions between biochemical and morphological predictors significantly affected TE convexity, but were not omitted in the main figures for clarity. The lines represent the effect of the respective variable on convexity when the interactor value was high (solid line), average (dashed line), or low (dotted line).

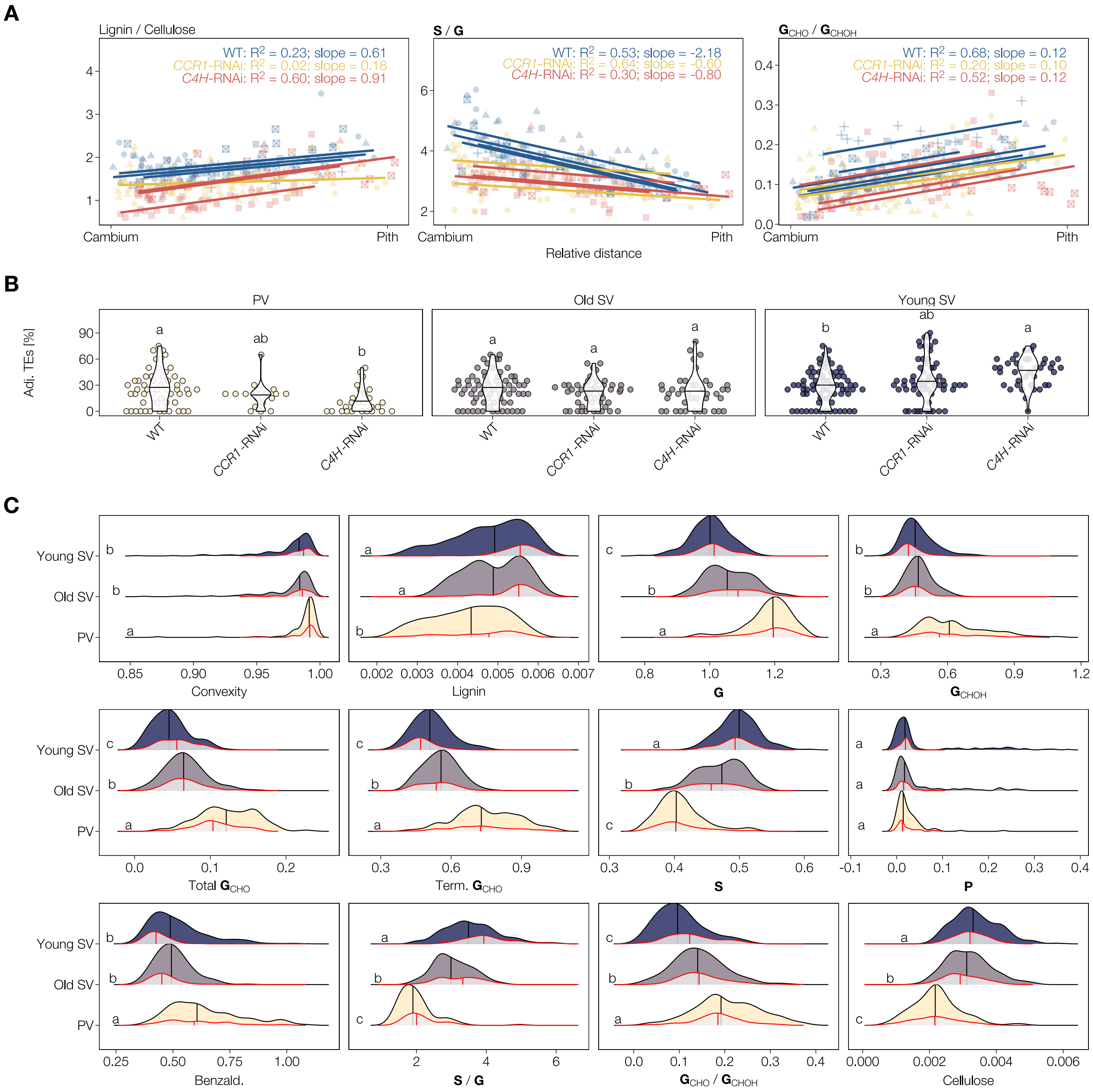


**Figure S6 | TE morphology and lignin composition in *Populus tremula* × *tremuloides* phenylpropanoid mutants (related to figure 5).** **A** *Post-mortem* lignification in *Populus tremula* × *tremuloides* TEs. The ratios of lignin to cellulose, **S** to **G**, and $\mathbf{G}_{\mathrm{CHO}}$ to $\mathbf{G}_{\mathrm{CHOH}}$ in secondary TEs of poplar stems changed with their distance to the cambium, *i.e.* their age. The lines, conditional $R^2$ values and slopes represent mixed linear models of the respective ratio against the distance from the cambium, allowing for different intercepts for each plant. **B** Relative proportion of TE perimeters surrounded by other TEs in the different genotypes comprising our dataset. Letters indicate significant differences between genotypes according to a Tukey-HSD test (per panel; $\alpha$ = 0.05). **C** *Populus tremula* × *tremuloides* TE convexity and cell wall composition data used in the structural equation models (Fig. 5 H–J). Variation across all genotypes (in blue/purple/yellow) overlayed with the variation in the WT (grey with red outline, scaled to 30%). Vertical lines represent the respective median values. Letters indicate significant differences between TE morphotypes according to a Kruskal-Wallis test followed by Dunn's multiple comparison (per panel; $\alpha$ = 0.05). All data used in the models is also available in the supplemental data file.

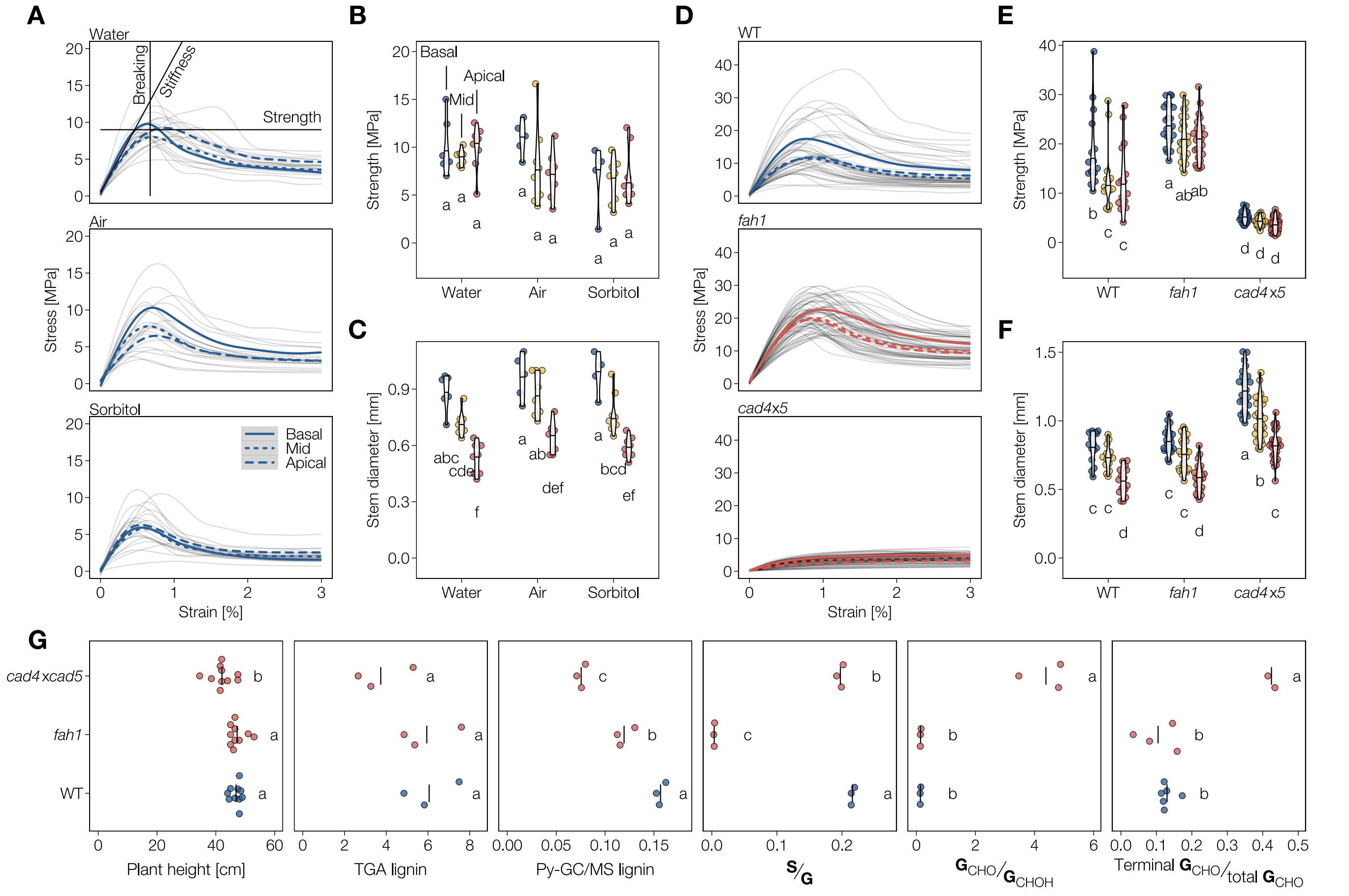


**Figure S7 | Stem biomechanics and lignin composition in *fah1* and *cad4×5* (related to figure 6). A** Bending curves of the three-point-bending test of WT plants incubated in different media. Average bending curves for each developmental stage (basal, mid, apical) are indicated by blue lines. The strength is defined as the maximum endured stress (solid line), whereas stiffness is calculated from the linear phase of the elastic bending (dashed line). The point of breaking is defined as the amount of strain the stem can endure before catastrophic failure, and is visible as the peak of the bending curve (dotted line). **B** Flexural strength of WT stems tested after incubation in different media. **C** Diameters of WT stems used in the bending experiments assessing the influence of different media. **D** Bending curves of mutant plant stems. **E** Flexural strength of mutant plant stems. **F** Diameters of tested mutant plant stems. **G** Plant height, total lignin according to thioglycolic acid derivatisation (TGA) and pyrolysis-GC/MS (Py-GC/MS), **S**/**G** and $\mathbf{G}_{CHO}$/$\mathbf{G}_{CHOH}$ ratios according to Py-GC/MS, and terminal $\mathbf{G}_{CHO}$/$\mathbf{G}_{CHOH}$ ratio according to thioacidolysis-GC/MS. Dots represent individual samples (single plants or pools of several plants), vertical lines represent the average for each variable and genotype.
Letters indicate significant differences according to a Tukey-HSD test (per panel; $\alpha$ = 0.05).

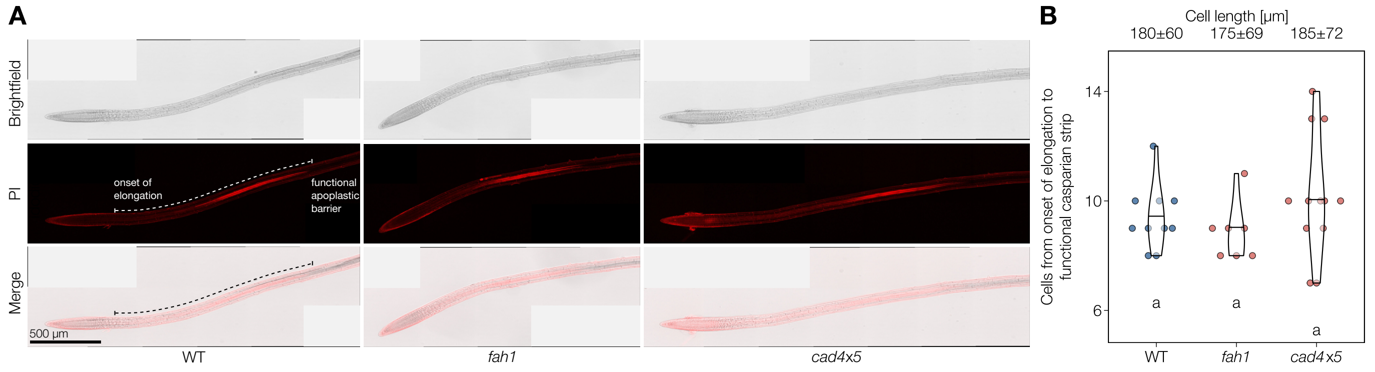


**Figure S8 | The Casparian strip is unaffected in *fah1* and *cad4×5* (related to figure 7). A** Root tips of four day old seedlings stained with propidium iodide (PI), indicating a functional apoplastic barrier in the endodermis excluding the dye from staining the vascular cylinder. **B** Establishment of the apoplastic barrier in $n$ = 7–12 individual seedlings occurred roughly 10 cells after the onset of elongation, unaffected by mutations in *F5H* (*fah1*) or *CAD4* and *5* (*cad4×5*) compared to WT plants. Letters indicate significant differences according to a Tukey-HSD test ($\alpha$ = 0.05).

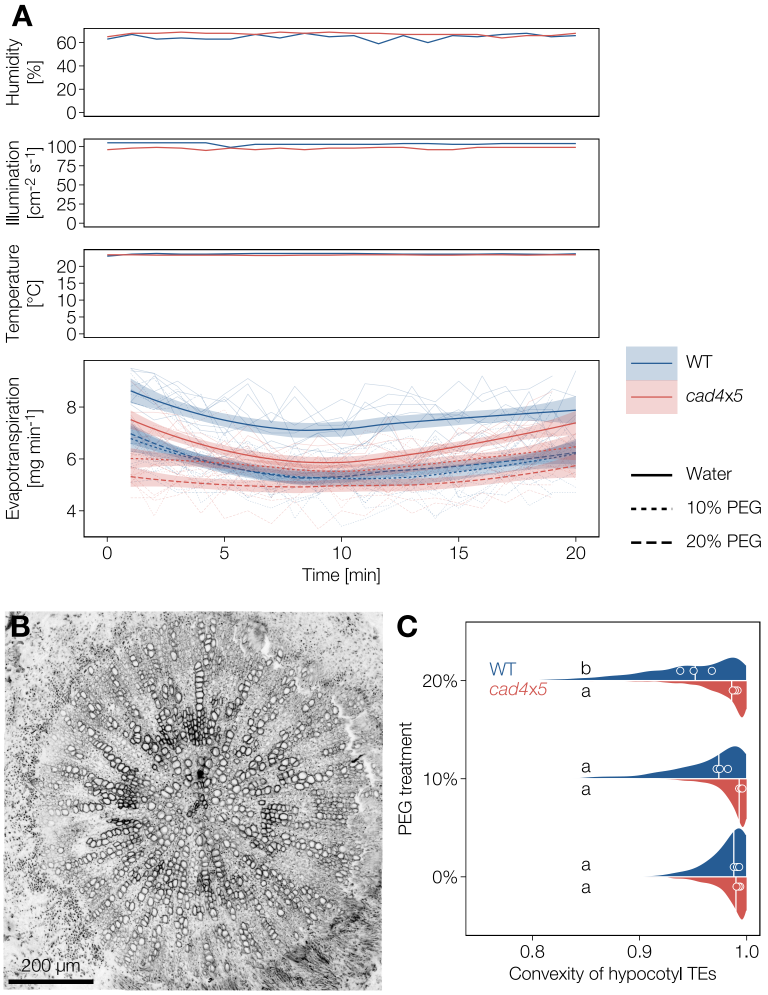


**Figure S9 | Overaccumulation of $G_{CHO}$ in *cad4×5* improves TE convexity after recovery from drought (related to figure 7). A** Variations in conditions during the evapotranspiration experiment. **B** Hypocotyl cross-section from a *cad4×5* plant after 10% PEG treatment and subsequent recovery in water. **C** TE collapse in hypocotyls after PEG treatment and subsequent recovery in water. The distribution and median lines represents all measured TEs, median convexity for each individual plant is indicated by points. Letters indicate significant differences between genotypes and treatments according to a Tukey-HSD test ($\alpha$ = 0.05); *n* = 3 individual plants per genotype and treatment.

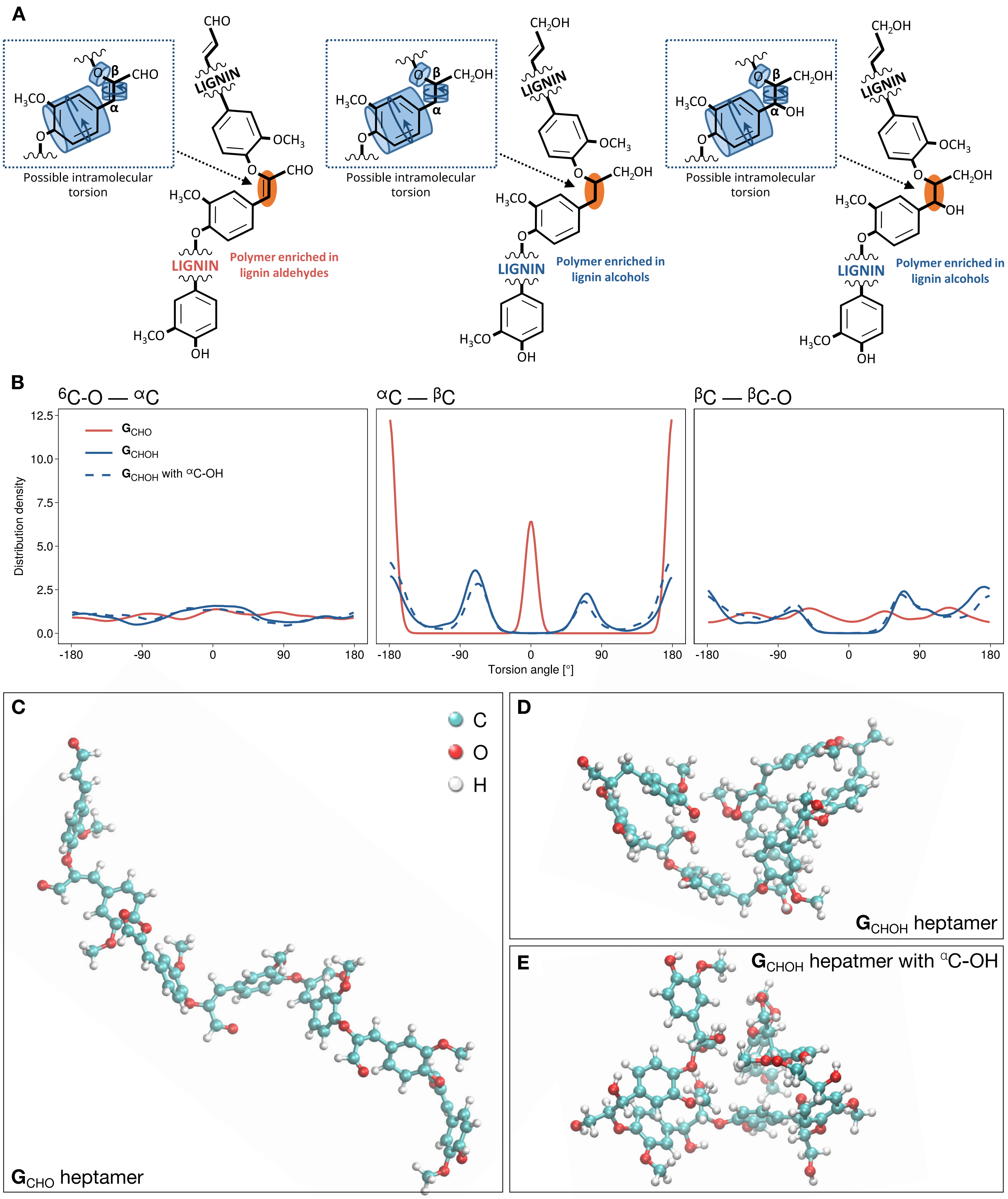


**Figure S10 | Topology and mechanics of lignin oligomers depends on the $C_3$ functional group (related to table 1). A** Schematic representation of the structure and torsion of ether linkage between residue of lignin oligomers depending on their aliphatic terminal function. **B** Distribution of torsion angles around the $^{6}$C-O—$^{\alpha}$C, the $^{\alpha}$C—$^{\beta}$C and the $^{\beta}$C—$^{\beta}$C-O atoms for lignin oligomers made of **G**$_{CHO}$ residues (red line) or **G**$_{CHOH}$ residues (blue line) without (solid line) or with $^{\alpha}$C-OH (dotted line) determined from molecular dynamics simulations. **C–E** Typical molecular conformations from equilibrated molecular dynamics simulations of lignin heptamers with **G**$_{CHO}$ residues (**C**), **G**$_{CHOH}$ residues without $^{\alpha}$C-OH (**D**), and **G**$_{CHOH}$ residues with $^{\alpha}$C-OH (**E**).

**Table S1 |** Used nomenclature of lignin chemistry.

| $C_6$ ring substitution | Aliphatic length | Aliphatic function | Abbreviation | Name |
|---|---|---|---|---|
| **G** | $C_3$ | –CHO | $\mathbf{G}_{CHO}$ | coniferaldehyde |
| **G** | $C_3$ | –CHOH | $\mathbf{G}_{CHOH}$ | coniferyl alcohol |
| **G** | $C_1$ | –CHO | benzaldehyde | vanillin |
| **G** | $C_3$ | –CHO/–CHOH/–COOH | **G** | guaiacyl residues |
| **S** | $C_3$ | –CHO/–CHOH/–COOH | **G** | syringyl residue |
| **S** | $C_1$ | –CHO | benzaldehyde | syringaldehyde |
| **P** | — | — | **P** | phenyl residues |

**Table S2 |** Insertional mutants and the targeted genes used in the present study including gene name, locus number, number of paralog for each plant species as well as previous references in which these plants were further analysed.

| Acronym | Locus | Full name | Plant species | Paralogs | Mutant alleles | Mutant ID | References |
|---|---|---|---|---|---|---|---|
| *C4H* | Potri.013G157900 | *cinnamate-4-hydroxylase* | hybrid poplar | 4 | | | Bjurhager et al. 2010 |
| *4CL1* | At1g51680 | *4-coumarate-CoA ligase 1* | Arabidopsis | 4 | *4cl1-1* | SALK_142526 | Van Acker et al. 2013 |
| *4CL2* | At3g21240 | *4-coumarate-CoA ligase 2* | Arabidopsis | 4 | *4cl2-4* | SALK_110197 | Li et al. 2015 |
| | | | Arabidopsis | 4 | *4cl1-1*×*4cl2-4* | | Blaschek et al. 2020 |
| *CCoAOMT1* | At4g34050 | *caffeoyl-CoA O-methyltransferase 1* | Arabidopsis | 1 | *ccoaomt1* | SALK_151507 | Kai et al. 2008 |
| *F5H1* | At4g36220 | *ferulate-5-hydroxylase 1* | Arabidopsis | 2 | *fah1-2* | EMS mutant | Meyer et al. 1998 |
| *OMT1* | At5g54160 | *caffeic acid O-methyltransferase 1* | Arabidopsis | 1 | *omt1* | SALK_135290 | Tohge et al. 2007 |
| *CCR1* | At1g15950 | *cinnamoyl-CoA reductase* | Arabidopsis | 2 | *ccr1-3* | SALK_123-689 | Mir Derikvand et al. 2008 |
| *CCR* | Potri.003G181400 | *cinnamoyl-CoA reductase* | hybrid poplar | 15 | | | Escamez et al. 2017 |
| *CAD4* | At4g37980 | *cinnamyl alcohol dehydrogenase 4* | Arabidopsis | 9 | *cad4-1* | SAIL_1265_A06 | Lee et al. 2017 |
| *CAD5* | At4g37990 | *cinnamyl alcohol dehydrogenase 5* | Arabidopsis | 9 | *cad5-1* | SAIL_776_B06 | Lee et al. 2017 |
| | | | Arabidopsis | 9 | *cad4-1*×*cad5-1* | | Blaschek et al. 2020 |

Paralog numbers were taken from Raes et al. (2003) for *Arabidopsis thaliana* and from Sundell et al. (2017) for hybrid poplar

**Table S3 |** Test statistics on global goodness-of-fit (Fisher's C) and directed separation (*i.e.* independence of variables) in the piecewise structural equation models (related to figures 4 and 5).

| Species | Cell type | Fisher's C | *P*-value[a] | Independence claim | Crit. value | *P*-value[b] |
|---|---|---|---|---|---|---|
| *Arabidopsis* | PX | 16.650 | 0.034 | Circularity ~ Perimeter + ... | -2.4405 | 0.0163 |
| | | | | Circularity ~ $\mathbf{G}_{CHOH}$ + ... | 1.6967 | 0.0926 |
| | | | | Circularity ~ Terminal $\mathbf{G}_{CHO}$ + ... | -0.2495 | 0.8034 |
| | | | | Circularity ~ Total lignin + ... | 1.2890 | 0.2001 |
| | MX | 23.020 | 0.003 | Circularity ~ Total $\mathbf{G}_{CHO}$ + ... | 1.0932 | 0.2769 |
| | | | | Circularity ~ **S** + ... | -2.4661 | 0.0153 |
| | | | | Circularity ~ Terminal $\mathbf{G}_{CHO}$ + ... | 1.7678 | 0.0801 |
| | | | | Circularity ~ Total lignin + ... | 2.2066 | 0.0296 |
| | SX | 6.996 | 0.136 | Circularity ~ $\mathbf{G}_{CHOH}$ + ... | 0.2350 | 0.8148 |
| | | | | Circularity ~ Benzaldehydes + ... | 2.1184 | 0.0371 |
| Poplar | Old SV | 4.942 | 0.764 | Circularity ~ **P** + ... | 0.9855 | 0.3259 |
| | | | | Circularity ~ $\mathbf{G}_{CHOH}$ + ... | 0.1769 | 0.8598 |
| | | | | Circularity ~ Terminal $\mathbf{G}_{CHO}$ + ... | 0.8703 | 0.3855 |
| | | | | Circularity ~ Total lignin + ... | 0.2767 | 0.7824 |
| | Young SV | 13.135 | 0.216 | Circularity ~ Perimeter + ... | 2.2784 | 0.0240 |
| | | | | Circularity ~ **P** + ... | 0.8344 | 0.4052 |
| | | | | Circularity ~ $\mathbf{G}_{CHOH}$ + ... | -0.2598 | 0.7954 |
| | | | | Circularity ~ **S** + ... | -1.2235 | 0.2229 |
| | | | | Circularity ~ Terminal $\mathbf{G}_{CHO}$ + ... | -0.2332 | 0.8159 |

PX, protoxylem TEs; MX, metaxylem TEs; SX secondary xylem TEs; PV, primary vessels; SV, secondary vessels

Independence claims are formatted as (response variable) ~ (predictor variable) + ... (*i.e.* within the models defined in Figs. 4 and 5)

[a] From the global goodness-of-fit test; values < 0.05 indicate that a theoretically better model can be built by including additional relationships between parameters.

[b] From the test of directed separation; values < 0.05 indicate that the respective relationship is statistically significant.

**Table S4 |** Standardised and raw coefficients and their *P*-values in the piecewise structural equation models (related to figures 4 and 5).

| Species | Cell type | Response | Predictor | Estimate | Std. error | Crit. value | *P*-value | Std. estimate[a] |
|---|---|---|---|---|---|---|---|---|
| *Arabidopsis* | PX | Circularity | Convexity | 1.9018 | 0.0633 | 30.0486 | 0.0000 | 0.9441 |
| | | Convexity | Perimeter | 0.0078 | 0.0012 | 6.5366 | 0.0000 | 0.4690 |
| | | Convexity | $\mathbf{G}_{CHOH}$ | 0.1949 | 0.0536 | 3.6377 | 0.0004 | 0.2674 |
| | | Convexity | Terminal $\mathbf{G}_{CHO}$ | 0.1228 | 0.0435 | 2.8237 | 0.0057 | 0.2041 |
| | | Convexity | Total lignin | 19.5650 | 6.8561 | 2.8537 | 0.0052 | 0.2045 |
| | MX | Circularity | Adjacent vessels | -0.0537 | 0.0274 | -1.9615 | 0.0525 | -0.0677 |
| | | Circularity | Convexity | 2.0009 | 0.0748 | 26.7501 | 0.0000 | 0.9231 |
| | | Convexity | Adjacent vessels | -0.0626 | 0.0270 | -2.3181 | 0.0225 | -0.1712 |
| | | Convexity | Total $\mathbf{G}_{CHO}$ | 0.2336 | 0.0891 | 2.6216 | 0.0101 | 0.2172 |
| | | Convexity | **S** | -0.3491 | 0.1299 | -2.6872 | 0.0084 | -0.2950 |
| | | Convexity | Terminal $\mathbf{G}_{CHO}$ | 0.1427 | 0.0610 | 2.3416 | 0.0212 | 0.1902 |
| | | Convexity | Total lignin | 17.0394 | 8.1287 | 2.0962 | 0.0386 | 0.2237 |
| | SX | Circularity | Convexity | 2.0475 | 0.1507 | 13.5842 | 0.0000 | 0.8290 |
| | | Convexity | $\mathbf{G}_{CHOH}$ | 0.0660 | 0.0264 | 2.5042 | 0.0142 | 0.2221 |
| | | Convexity | Benzaldehydes | -0.1178 | 0.0195 | -6.0314 | 0.0000 | -0.5349 |
| Poplar | PV | Circularity | Adjacent vessels | -0.0111 | 0.0208 | -0.5327 | 0.5955 | -0.0306 |
| | | Circularity | Convexity | 3.6455 | 0.2520 | 14.4680 | 0.0000 | 0.8312 |
| | | Convexity | Adjacent vessels | -0.0164 | 0.0083 | -1.9707 | 0.0517 | -0.1982 |
| | Old SV | Circularity | Adjacent vessels | -0.0741 | 0.0141 | -5.2637 | 0.0000 | -0.2209 |
| | | Circularity | Convexity | 2.0701 | 0.1097 | 18.8682 | 0.0000 | 0.7918 |
| | | Convexity | Adjacent vessels | -0.0218 | 0.0079 | -2.7535 | 0.0066 | -0.1696 |
| | | Convexity | **P** | -0.1817 | 0.0247 | -7.3561 | 0.0000 | -0.5321 |
| | | Convexity | $\mathbf{G}_{CHOH}$ | 0.1282 | 0.0505 | 2.5380 | 0.0121 | 0.3109 |
| | | Convexity | Terminal $\mathbf{G}_{CHO}$ | -0.0795 | 0.0360 | -2.2090 | 0.0287 | -0.2481 |
| | | Convexity | Total lignin | 6.6005 | 2.4852 | 2.6559 | 0.0087 | 0.2019 |
| | Young SV | Circularity | Adjacent vessels | -0.1153 | 0.0131 | -8.8058 | 0.0000 | -0.3740 |
| | | Circularity | Convexity | 1.9740 | 0.1256 | 15.7185 | 0.0000 | 0.6675 |
| | | Convexity | Adjacent vessels | -0.0180 | 0.0064 | -2.8013 | 0.0057 | -0.1731 |
| | | Convexity | Perim. | -0.0002 | 0.0001 | -2.3985 | 0.0176 | -0.1620 |
| | | Convexity | **P** | -0.1195 | 0.0161 | -7.4287 | 0.0000 | -0.4652 |
| | | Convexity | $\mathbf{G}_{CHOH}$ | 0.2119 | 0.0404 | 5.2477 | 0.0000 | 0.6687 |
| | | Convexity | **S** | -0.3164 | 0.0490 | -6.4540 | 0.0000 | -0.4228 |
| | | Convexity | Terminal $\mathbf{G}_{CHO}$ | -0.2424 | 0.0374 | -6.4732 | 0.0000 | -0.8465 |

PX, protoxylem vessels; MX, primary metaxylem vessels; SX secondary metaxylem vessels; PV, primary vessels; SV, secondary vessels

[a] Standardised coefficients allow direct comparisons of relative impacts within each model, but not across different models.
